# Prioritisation of oncology therapeutic targets using CRISPR-Cas9 screening

**DOI:** 10.1101/502005

**Authors:** Fiona M Behan, Francesco Iorio, Emanuel Gonçalves, Gabriele Picco, Charlotte M Beaver, Rita Santos, Yanhua Rao, Rizwan Ansari, Sarah Harper, David Adam Jackson, Rebecca McRae, Rachel Pooley, Piers Wilkinson, David Dow, Carolyn Buser-Doepner, Euan A. Stronach, Julio Saez-Rodriguez, Kosuke Yusa, Mathew J Garnett

**Affiliations:** Wellcome Sanger Institute, Wellcome Genome Campus, Cambridge CB10 1SA, UK; Open Targets, Wellcome Genome Campus, Cambridge CB10 1SA, UK; European Molecular Biology Laboratory – European Bioinformatics Institute, Wellcome Genome Campus, Cambridge CB10 1SD, UK; GSK, Target Sciences, Gunnels Wood Road, Stevenage, SG1 2NY, UK and Collegeville, Pennsylvania, 19426–0989, USA; Faculty of Medicine, Joint Research Centre for Computational Biomedicine, RWTH Aachen University, Aachen 52057, Germany; Heidelberg University, Heidelberg, Germany

**Keywords:** Cancer, CRISPR, synthetic-lethal, screen, Werner helicase, therapeutics, cell lines, targets

## Abstract

Functional genomics approaches can overcome current limitations that hamper oncology drug development such as lack of robust target identification and clinical efficacy. Here we performed genome-scale CRISPR-Cas9 screens in 204 human cancer cell lines from 12 cancer-types and developed a data-driven framework to prioritise cancer therapeutic candidates. We integrated gene cell fitness effects with genomic biomarkers and target tractability for drug development to systematically prioritise new oncology targets in defined tissues and genotypes. Furthermore, we took one of our most promising dependencies, Werner syndrome RecQ helicase, and verified it as a candidate target for tumours with microsatellite instability. Our analysis provides a comprehensive resource of cancer dependencies, a framework to prioritise oncology targets, and nominates specific new candidates. The principles described in this study can transform the initial stages of the drug development process contributing to a new, diverse and more effective portfolio of oncology targets.

## Main Text

Cancer is the second leading cause of death globally and incidence rates are rising ^1^. The molecular features of a patient’s tumour impact the clinical responses to therapy and can be used to guide therapies, leading to more effective treatments and reduced toxicity ^2^. A paradigm in targeted cancer therapy is the use of drugs which target a dominant gain-of-function oncogene or activated downstream signalling pathway. Alternatively, therapies can exploit the stress phenotype induced by molecular alterations, such as loss of a tumour suppressor gene (e.g. *BRCA1* and *BRCA2*), which can create dependencies on other cellular pathways ^3^. Despite the increasing numbers of available targeted cancer therapies, most patients do not benefit from such therapies mainly due to the lack of knowledge on targetable alterations ^4,5^. The attrition rate in oncology drug development is ~90%, with a leading cause of failure being the lack of efficacy, and fewer molecular entities to new targets are being developed ^6^. Current approaches to target selection are frequently hampered by incomplete information, are subject to confirmation bias, and can be biased towards well studied pathways. Strategies that improve our ability to identify and prioritise drug targets could expand the number of targets, increase success rates and accelerate development of new cancer therapies.

Loss-of-function genetic screens are a powerful approach to comprehensively investigate gene function. RNA interference screens are useful ^7,8^, but have been hampered by high variation, off-target activity and incomplete knockdown. In contrast, the CRISPR-Cas9 system has greater specificity and produces more penetrant phenotypes resulting from null alleles generated ^9,10^. Libraries of single guide RNAs (sgRNAs) have been used in pooled genome-wide screens to study gene function and their requirement for cellular fitness ^11,12^. Although CRISPR-Cas9 screens have been reported, no study has performed a systematic integration of gene fitness effects, tractability for pharmaceutical development, and biomarkers for patient selection across a diverse panel of cancer cell lines to prioritise new therapeutic targets. Here, we present genome-scale CRISPR-Cas9 fitness screens in 204 cancer cell lines and a computational, data-driven analysis to prioritise cancer therapeutic targets, illustrated with the identification and validation of Werner syndrome helicase as a target for tumours with microsatellite instability.

### Genome-scale CRISPR-Cas9 screens in cancer cell lines

To comprehensively catalogue genes required for cancer cell fitness (defined as genes required for cell growth or viability) in diverse histological and molecular sub-types, we performed genome-scale CRISPR-Cas9 screens targeting 18,009 genes in 204 cancer cell lines across 12 different tissues, including lung (n = 43), ovarian (n = 35), colorectal (n = 34), peripheral/central nervous system (n = 29), pancreatic (n = 24), breast (n = 24), and other (n = 14) (**Figs. 1a, b** and **S1a**). The vast majority of the cell lines (98%) are part of the Genomics of Drug Sensitivity in Cancer cell line panel ^13^, where single nucleotide variants (SNVs) and gene copy number variations (CNVs) are fully annotated (**Fig S1b**). The cell lines selected broadly reflect the molecular features of patient tumours ^13^, include the three most common forms of cancer (lung, colon and breast), and cancers of particular unmet clinical need (lung, ovarian and pancreas).

We observed high concordance between technical replicates in each cell line when considering raw sgRNA counts (**Fig. S1c**, median R = 0.81). A more stringent quality control (QC) assessment using gene-level log2 fold-change (logFC) retained 95% of the replicates (median correlation of 0.84)(**Fig. S1d-f**), leaving a final analysis set of 197 cell lines (**Table S1**). The analysis set showed high sensitivity, specificity and precision in classifying essential and non-essential gene sets ^14^ based on gene-level logFC ranks (**Fig. S1g**), and robust depletion of known essential gene sets (**Fig. 1c** and **S1h**; grand median logFC = −2.92 and −1.91, and median Glass’ Δ (GΔ) = 2.80 and 2.22 for ribosomal and the essential genes, respectively). These results demonstrate the high quality of our dataset and accuracy to detect fitness genes. We performed CRISPR-bias ^15,16^ correction on the 197 cell line dataset using *CRISPRcleanR* ^17^, and computed gene-level fitness scores using modified versions of BAGEL ^18^ and MAGeCK ^19^ to catalogue fitness genes.

**Figure 1:**
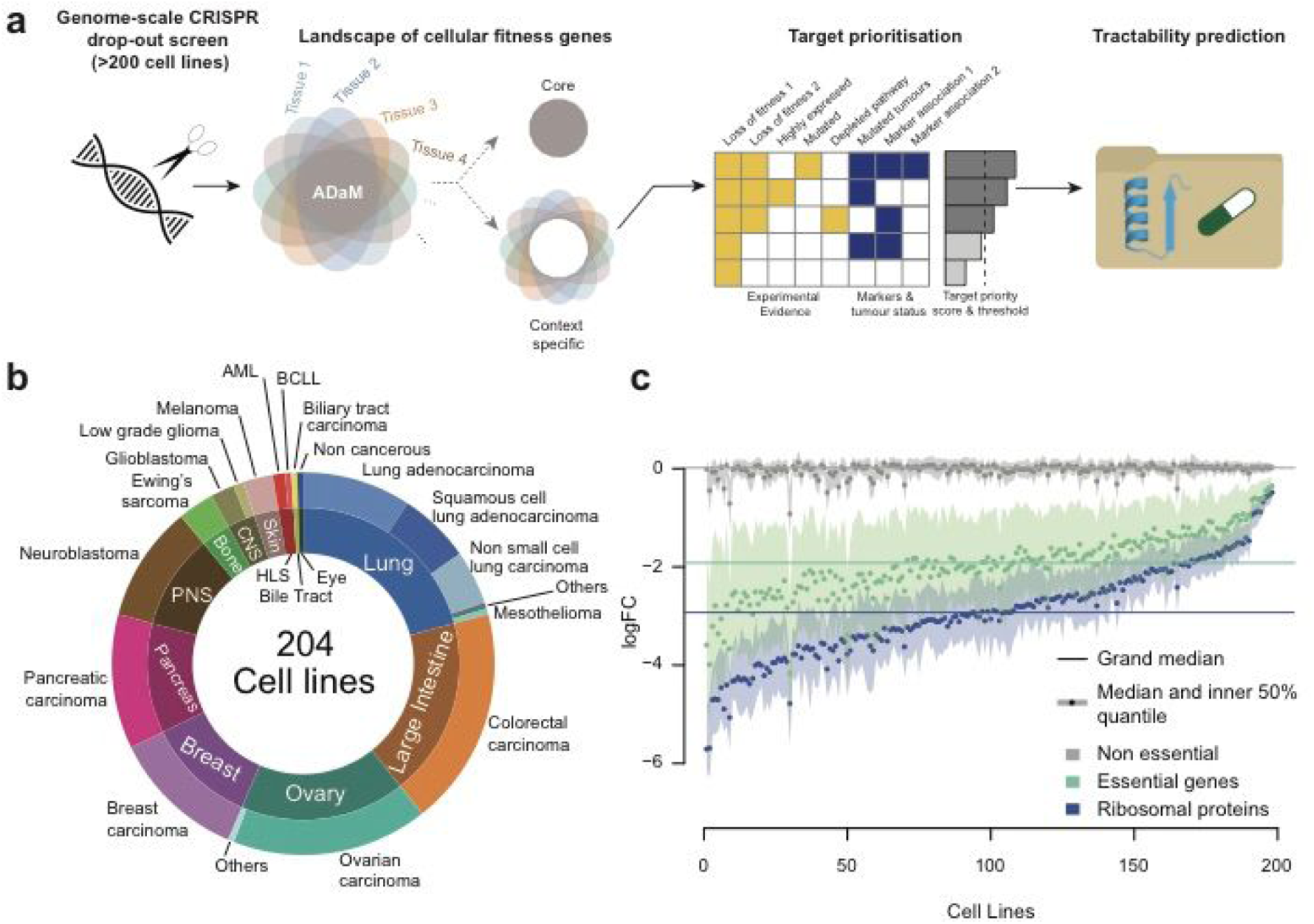
Genome-scale CRISPR-Cas9 screens in cancer cell lines. (a) Schematic of the target prioritisation strategy incorporating context specific gene fitness effects, genomic biomarker and target tractability information. (b) Cancer cell lines screened grouped by tissue (inner ring) and cancer-type (outer ring). (c) Median logFC values (averaged across targeting sgRNAs) and inter-quartiles for defined sets of genes across cell lines.

### Landscape of cellular fitness genes

We identified a median of 1,413 fitness genes across all cell lines (BAGEL FDR < 5%; **Figs. 2a, S2a**, and **Table S2**). Technical confounders (e.g. number of replicates and Cas9 activity) were not robustly associated with the number of fitness genes identified (**Fig. S2b-e**). In total, 38% of all targeted genes (n = 6,830) had a loss of fitness effect in one or more cell lines (hereafter referred to as vulnerable cell lines), in line with the ~42% identified in mice ^20^. The majority (82%) of fitness essential genes induced a vulnerability in less than 50% of cell lines (**Fig. 2b**). Genes required for fitness in specific contexts are more likely to make favourable drug targets because of reduced likelihood of toxicity in healthy tissues. Conversely, fitness genes common across all cancer-types (referred to as pan-cancer core fitness (CF) genes) or in a given cancer-type (cancer-type CF genes) are likely to have higher toxicity. It is therefore important to distinguish context-specific fitness genes from CF genes during target prioritisation. Utilising our large dataset, we developed a novel statistical method, ADaM (adaptive daisy model, **Fig. S3**), to identify CF genes of a given cell line set. ADaM adaptively determines the minimum number of vulnerable cell lines required for a gene to be classified as CF in a cell line set. This number maximises the true positive rates of known essential genes in the resulting CF set, and the deviation from expectation of the number of genes in this set. The ADaM approach estimated a median of 82% of cell lines from a cancer-type to be vulnerable for a gene to be considered as a cancer-type CF gene and at least 10 of 12 cancer-types for pan-cancer CF genes, yielding a median of 764 cancer-type CF genes and 582 pan-cancer CF genes (**Fig. 2c**, and **Table S3**). Of note, although the majority of context-specific fitness genes exhibited weaker effect than CF genes, a subset of context-specific fitness genes had an effect size similar to or even stronger than the CF genes, indicating strong context-specific vulnerabilities (**Fig. 2c**).

Of the pan-cancer CF genes, 419 were a subset of the BAGEL essential genes ^14^ or a more recent set of CF genes ^21^, and 127 encompassed genes in other essential processes (histones, ribosome, proteasome, spliceosome and RNA-polymerase) ^22,23^. Interestingly, 140 (24%) were newly classified as CF genes with our screen dataset and enriched in housekeeping pathways (mediator complex, anaphase promoting complex, and general transcription factors) and pathways involved in cell growth and division (sister chromatid separation and protein transcription) (**Fig. S4a** and **b** and **Table S4**). Compared to reference sets of CF genes (^14^, ^21^), the ADaM pan-cancer CF genes showed a greater recall of genes involved in essential processes (median = 67% versus 28% and 51% respectively, **Fig. S4c**), and a similar false discovery rates for putative context-specific fitness genes (taken from ^8^, **Fig. S4d**). Clustering of cancer-types based on corresponding CF gene sets similarity reflected their tissue of origin, with blood cancer cell lines having the most distinctive set of CF genes (342 exclusive CF genes; **Fig. S5a**). CF gene sets were highly expressed in matched healthy tissues (grand median expression > 75% quantile, p < 10^−16^; **Fig. S5b**), consistent with their predicted role in core cellular processes. Overall, this analysis enabled us to distinguish CF genes likely having greater toxicity when prioritising therapeutic targets, and expanded and refined our knowledge of human CF genes, likely having utility in many aspects of human genetics in health and disease.

**Figure 2:**
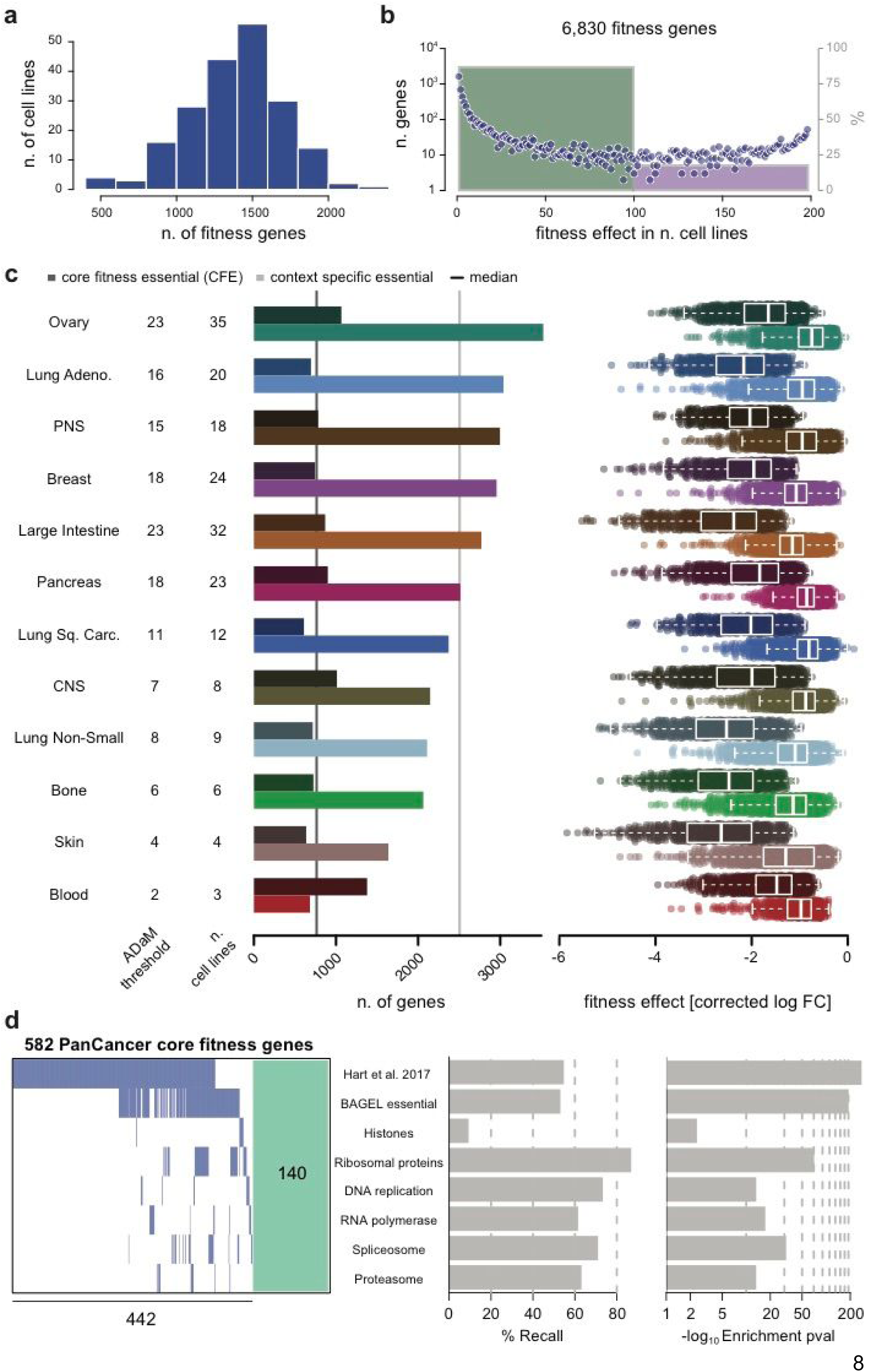
Landscape of fitness genes in a heterogeneous population of cancer cell lines. (a) The distribution of the number of fitness genes per cell line. (b) Number of genes exerting a fitness effect in a given number of cell lines. The bars show the percentage of genes which induce vulnerability in less (green bar) or more (purple bar) than 50% of cell lines. (c) Number of core fitness genes and context-specific fitness genes predicted by ADaM for each cancer-type and (on the right) average significant fitness effect for genes in the two sets (only statistically significant loss of fitness effects at a 5% BAGEL FDR are considered). The ADaM threshold is the number of cell lines a gene must be called as a fitness gene to be classified as core fitness essential. (d) Pan-cancer core fitness genes and their membership in known reference essential gene sets and respective recall and enrichment significance.

### A quantitative framework for target prioritization

Through our CRISPR screens we identified a large number of context-specific fitness genes. Nonetheless, it was unclear which genes, and what proportion, could represent potential drug targets. To nominate the most promising therapeutic targets, we developed a computational framework that integrates multiple lines of evidence and creates data-driven ranked lists of candidates at a pan-cancer and individual cancer-type level (**Fig. S6a**). We assigned each gene a target priority score between 0 and 100 (low to high). 70% of the score was derived from CRISPR-Cas9 experimental evidence and averaged across vulnerable cell lines based on (i) effect size, (ii) significance of fitness deficiency, (iii) basal gene expression, (iv) mutational status, and (v) evidence for other fitness genes in the same pathway. The remaining 30% was based on (i) evidence of an associated genetic biomarker and (ii) the frequency at which the target is somatically altered in patient tumours. For the biomarker analysis, we performed an analysis of variance (ANOVA, **Fig. S7**) to test associations between gene-level logFC of fitness genes and the presence of 381 cancer driver events (105 SNVs and 276 CNVs) ^13^ or microsatellite instability (MSI) status at pan-cancer and individual cancer-type levels (see Methods). To exclude genes likely to be poor targets due to potential toxicity core fitness genes were scored as zero, as well as potential false positive fitness genes (i.e. not expressed or homozygously deleted genes). Lastly, we defined a priority score threshold (52.6 and 45.23 for pan-cancer and cancer-type specific, respectively) based on scores calculated for targets with approved or pre-clinical cancer compounds (**Fig. S6b** and **Table S6**; see Methods). Priority targets were then classified based on a score greater than the defined thresholds.

With this approach, we identified 497 unique priority targets (20% of all unique targets with at least one non-null score, n = 2,537), including 83 and 470 targets from the pan-cancer and cancer-type-specific analyses, respectively (**Fig. 3a, Table S7**). The majority of the priority pan-cancer targets (67%) were also identified in cancer-type-specific analyses (**Fig. S6c**). The remaining 27 genes identified uniquely as pan-cancer priority targets typically induced vulnerability in a small subset of cell lines across multiple cancer-types (e.g. *NRAS* and *WWTR1*), or in a cancer-type where limited numbers of cell lines were available, thereby being precluded in the cancer-type specific analysis (*FLI1* in Ewing’s sarcoma; **Fig. S6d**). The number of priority targets varied ~2-fold across cancer-types, with a median of 133 targets per cancer-type (**Fig. 3a**). Significantly, the majority of cancer-type priority targets (n = 382, 81%) were detected in only one (59%) or two cancer-types (22%), underscoring the tissue specificity of many therapeutic targets.

Of the 497 priority targets, 59 (12%) were found to be robustly associated with at least one cancer driver event, and high significance and large effect size in fitness deficiency, and thus would be of particular interest for drug development (**Fig. 3b**). These targets (named Class A targets) included 29 pan-cancer and 34 cancer-type priority targets. Applying a less stringent threshold expanded the set of targets with cancer driver event associations (thus defining Class B and C targets), some of which were identified in multiple cancer-types (**Table S8**). For example, *PIK3CA* is a Class A target in breast, lung, colorectal and ovarian carcinoma, and PI3K inhibitors are in clinical development for *PIK3CA* mutated cancers ^24^. Taken together, these results highlight the potential of a quantitative framework, aggregating CRISPR-Cas9 screening data across multiple cell lines with associated genomic features, in prioritizing oncology targets.

**Figure 3:**
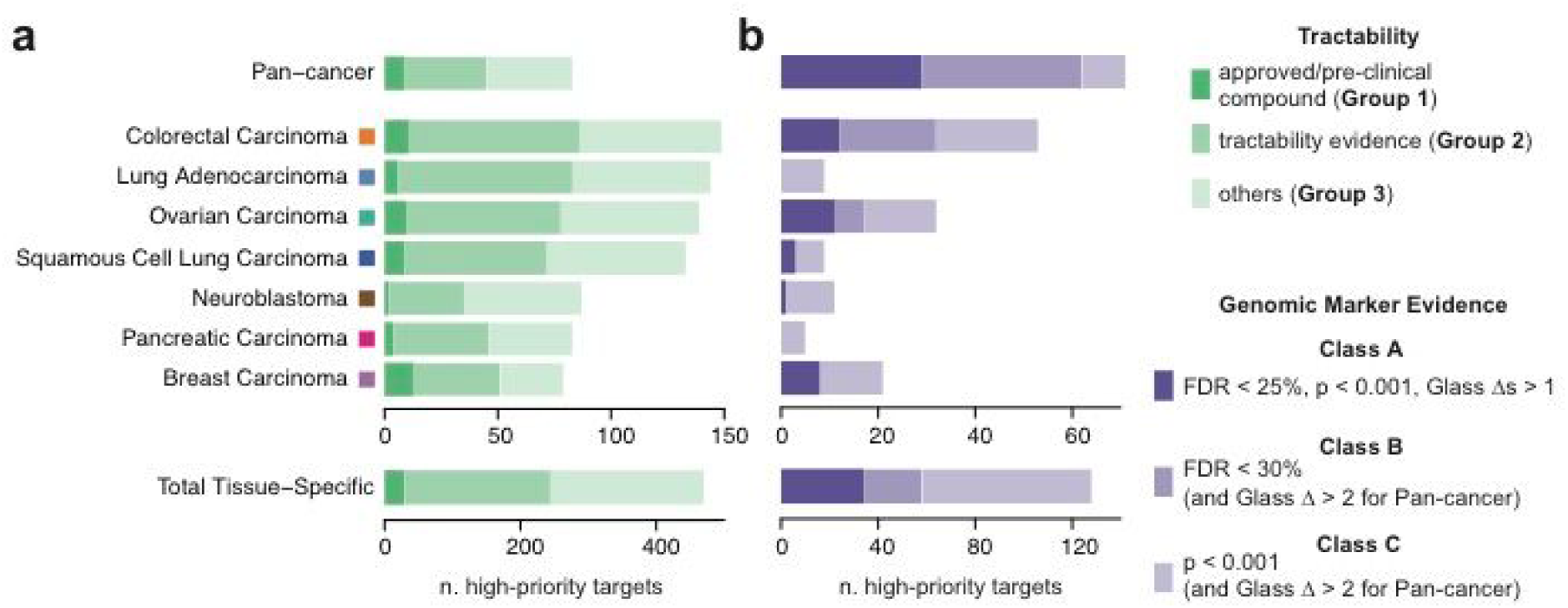
Target prioritization in different cancer-types. (a) The number of priority targets for each cancer-type and their tractability for drug development indicated by shading. (b) The number of priority targets with a genomic marker association. The shading indicates the statistical significance and effect size of the association.

### Tractability assessment of priority targets

Targets vary in their suitability for pharmaceutical intervention and this informs target selection, particularly during early stages of drug development. We previously conducted target tractability assessment for small molecule and antibody development, and assigned each gene to one of 10 tractability buckets (1–10 being high-to-low tractability) ^25^. This assessment uses multiple publicly available resources and considers factors such as existing (pre-)clinical compounds, patent literature supporting the target, availability of structural information, evidence of small molecule binding, or antibody accessibility. We cross-compared the 497 priority targets with their tractability and categorised them into three tractability groups (**Figs. 3b** and **4** and **Table S9**).

Tractability Group 1 (buckets 1–3) are targets with approved anti-cancer drugs or drugs in clinical development, and included 32 priority targets (3 pan-cancer and 29 cancer-type specific), including *ERBB2, ERBB3, CDK4, AKT1, ESR1*, *TYMS* and *PIK3CB* in breast carcinoma, and *EGFR, PIK3CA*, *IGF1R*, *MTOR*, and *ATR* in colorectal carcinoma (**Fig. 4**). Thirteen Group 1 targets have at least one drug developed for the cancer-type in which the target was identified as priority. The remaining 22 targets have drugs that have been used or developed for treatment of other cancer-types, thereby representing opportunities for drug repurposing.

Of the 32 Tractability Group 1 targets, 16% have a Class A biomarker (i.e. they are associated with a large-effect size to a cancer driver event), indicating highly desirable targets (**Figs. 3b**). An example is *CSNK2A1*, which is a highly significant fitness gene in colorectal cancer cell lines with gain of a segment containing *FLT3* and *WASF3* (*p* = 2.8 × 10^−6^, GΔ > 2.8, **Fig. 5a**) and targeted by Silmasertib, a drug in development for haematological malignancies and solid tumours. Other examples of Group 1 targets with class A - C markers are: *ERBB2* dependency in the presence of *ERBB2* amplification; *BRAF* dependency in the presence of *BRAF* mutation; *PIK3CA* dependency in the presence of *PIK3CA* mutations; *PIK3CB* dependency in breast cancers cell lines with *PTEN* mutations, and *CDK4* in *CCND1-amplified* cells (**Fig. 5a** and **Table S5**). Tractability Group 1 targets were enriched in protein kinases (adjusted *p* = 1.22 × 10^−7^), highlighting a major focus in drug development, compared to Groups 2 and 3 (adjusted *p* = 3.33 × 10^−2^ and > 0.05, respectively), which include a more functionally diverse set of targets (**Fig. S8a**, and **Table S10**).

Tractability Group 2 (buckets 4–7) contained 222 priority targets, for which no drug is currently in clinical development but there is evidence supporting their tractability (**Fig. 3b** and **4**). Of these, 10% have a Class A biomarker making them attractive for drug development (**Fig 3b**). Notable examples with biomarkers include: *KRAS* in *KRAS* mutant cancer cell lines; *USP7* in *APC* wild-type colorectal cell lines; *KMT2D* in breast cancer cell lines with amplification of a chromosomal segment containing *PPM1D* and *CLTC*; *BCL2L1* in squamous cell lung carcinoma cell lines without loss of *ERCC3;* and *WRN* in MSI-H cell lines (**Fig. 5b**). Of the Group 2 targets that were not associated with a biomarker, *GPX4* was a priority target in multiple cancer-types such as breast, ovarian, pancreatic, and squamous cell lung cancers (**Fig 4**). Sensitivity to GPX4 inhibition has been recently shown to be associated with epithelial-mesenchymal transition (EMT) ^26,27^ and we observed differential EMT marker expression between GPX4-sensitive and insensitive cell lines (**Fig. S8b** and **Table S11**). This example is indicative of future refinements of our target prioritisation scheme to capture priority targets associated with an expanded set of molecular features including gene expression, chromatin and differentiation states.

Lastly, Group 3 included 243 priority targets with no support or a lack of information to inform on target tractability (**Fig. 3b** and **4**). These are overrepresented by transcription factors (adjusted *p* = 6.83 × 10^−7^; **Fig. S8a** and **Table S10**), such as *MYCN* in neuroblastoma, *FOSL1* in pancreatic carcinoma, and *GATA3, TFDP1* and *FOXA1* (*FOXA1* is associated with a Class A biomarker; **Fig. 5c**) in breast carcinoma, which are largely intractable for drug development.

**Figure 4:**
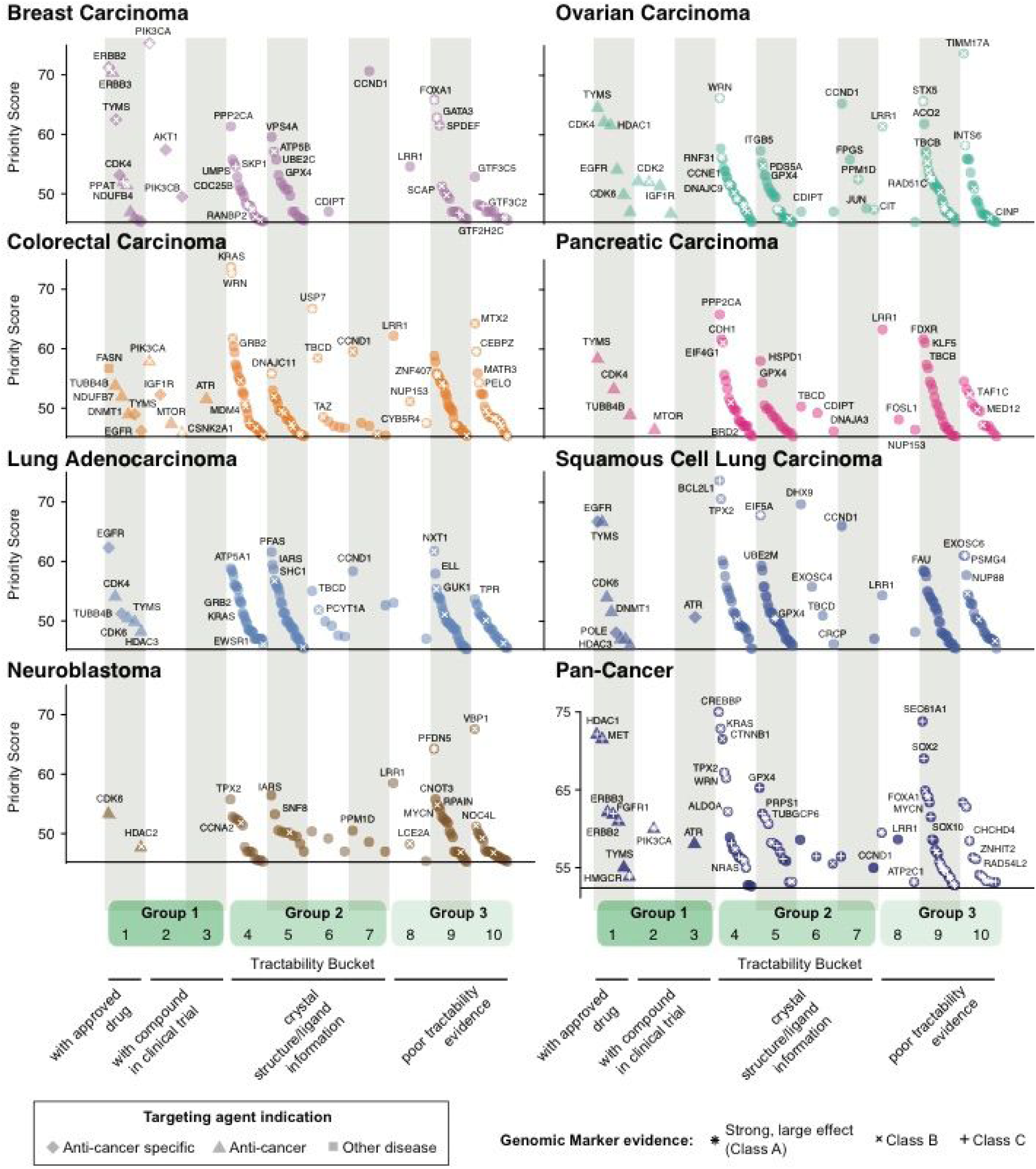
Priority therapeutic targets. Priority targets by cancer-type distributed across tractability buckets and groups. Each point is a target with a priority score (y-axis) in the indicated cancer-type or pan-cancer analysis. Shapes represent the indication of the approved/pre-clinical compound targeting the corresponding gene, circles indicate the absence of a compound. Symbols within the shape indicate the strength of the evidence of a biomarker associated with a differential essentiality of the target (respectively class A to C, from strong to weak evidence).

Priority targets were observed in all tractability groups, including a functionally diverse set of targets in Groups 2 and 3 (**Fig. 3b**). Targets in Group 2 are most likely to be novel and tractable to conventional modalities, and therefore represent good candidates for future drug development. Newer therapeutic modalities, such as proteolysis targeting chimeras (PROTACs, ^28^), and cell and gene therapy, may widen the range of proteins amenable to pharmaceutical intervention, especially those in Group 3 priority targets. Overall, our framework informed a data-driven list of prioritized therapeutic targets serving as strong candidates for oncology drug development.

**Figure 5:**
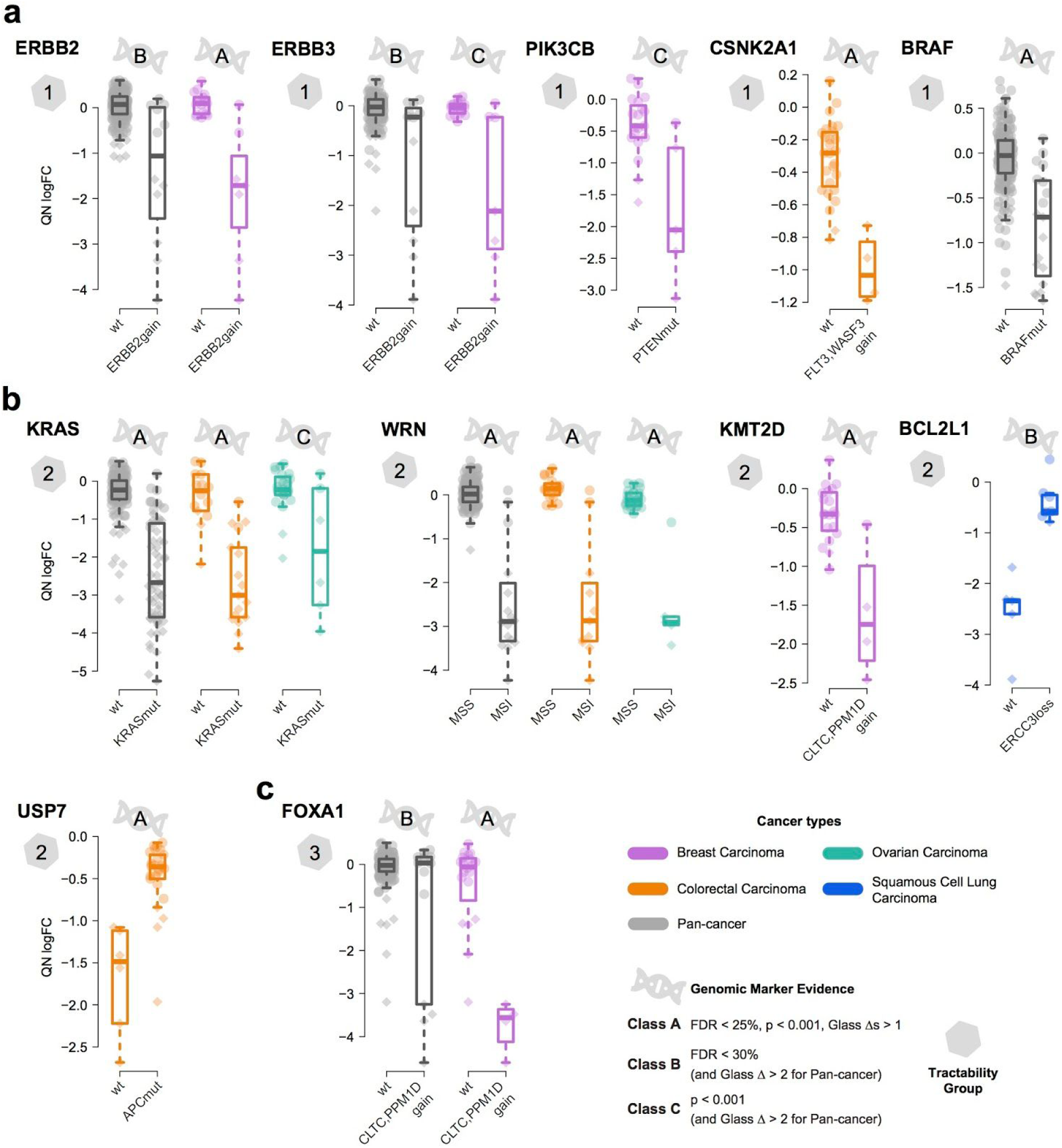
Priority target associated with a biomarker. **(a - c)** Differential fitness effect (quantile normalised depletion logFCs) for selected priority targets comparing cells with or without the associated genomic marker. Targets are grouped by tractability. An indicator of genomic marker class and tractability group is shown for each priority target. For some targets the same biomarker was identified within the pan-cancer and cancer-type specific analyses. Each point is a cell line, with color representing tissue type and shape indicating loss of fitness effect significance (with diamond indicating a BAGEL false discovery rate < 5%). Box and whiskers are interquartile ranges and 95th percentiles.

### Werner syndrome helicase is a target in cancers with microsatellite instability

Amongst the Class A priority targets, we identified the *WRN* gene encoding Werner syndrome RecQ like helicase as a promising candidate in colon and ovarian cancer cell lines (**Fig. 4**). Members of the RecQ DNA helicase family play roles in DNA repair, replication, transcription and telomere maintenance ^29^. WRN contains a helicase and endonuclease domain but currently has no clinical inhibitors available, and thus falls in Tractability Group 2 (Bucket 4). Vulnerability to *WRN* knockout was associated with high microsatellite instability (MSI-H) (ANOVA FDR = 1.38 × 10^−50^; **Figs. 5b** and **S7d**), as well as *ARID1B* mutations (ANOVA FDR = 6.82 × 10^−3^, **Table S5**); ARID gene family members have been previously linked with MSI ^30^. In contrast, other RecQ family members (*BLM, RECQL* and *RECQL5*) were not detected as fitness genes in the MSI-H cell lines. We mined data from systematic RNAi screens and confirmed preferential *WRN* vulnerability in MSI-H cancer cell lines (p-value = 0.004, **Fig. S9a**) ^8^, providing validation in an independent orthogonal experimental system. MSI-H phenotype is caused by impaired DNA mismatch repair (MMR) due to silencing or inactivation of MMR pathway genes (*MLH1, MLH3, MSH2, MSH3, MSH6, PMS1* and PMS2), is associated with high mutational load, and occurs in more than 20 different tumours types including colon, ovarian, endometrial and gastric cancers ^31,32^. A focused analysis of non-synonymous mutations, promoter methylation and homozygous deletions on MMR pathway genes confirmed the significant association between *MLH1* promoter hypermethylation and *WRN* essentiality (ANOVA FDR = 1.38 × 10^−50^, **Fig. 6a**).

To validate dependency on *WRN* in MSI-H cancers, we performed a CRISPR-based co-competition assay comparing the fitness of WRN-depleted versus wild-type cells. Using 4 individual sgRNAs targeting *WRN* (2 from the original library and 2 independently designed, **Table S12**) we observed a decrease in the ratio of WRN-depleted compared to wild-type cells exclusively in MSI-H cell lines (**Figs. 6b** and **S9b**). This was verified for 3 MSI-H cell lines included in the original screen and supported further by 4 additional MSI-H cell lines representing gastric and endometrial carcinoma (**Table S13**). In contrast, there was no change in the ratio of WRN-depleted compared to wild-type cells in all microsatellite stable (MSS) cell lines from 4 different tissue types (**Figs. 6b** and **S9b**). Similarly, *WRN* was selectively essential for MSI-H cells in clonogenic assays (**Figs. 6c** and **S9c**). Western blotting confirmed WRN protein knockdown for all 4 sgRNAs used (**Fig. S9d**). Of note, the effect size of *WRN* depletion on cellular fitness was similar to CF genes in both the primary screen and validation experiments, suggesting a potent effect on cellular fitness (**Fig 6a** and **b**). Furthermore, the fitness effect was selective for *WRN* as transgene expression of mouse *Wrn*, which is resistant to the sgRNA used, rescued *WRN* dependency in MSI-H cells (**Fig. 6d**). These results establish for the first time a link between MMR-deficiency and dependency on WRN, and support WRN as a priority candidate target in MSI-H cancers.

**Figure 6:**
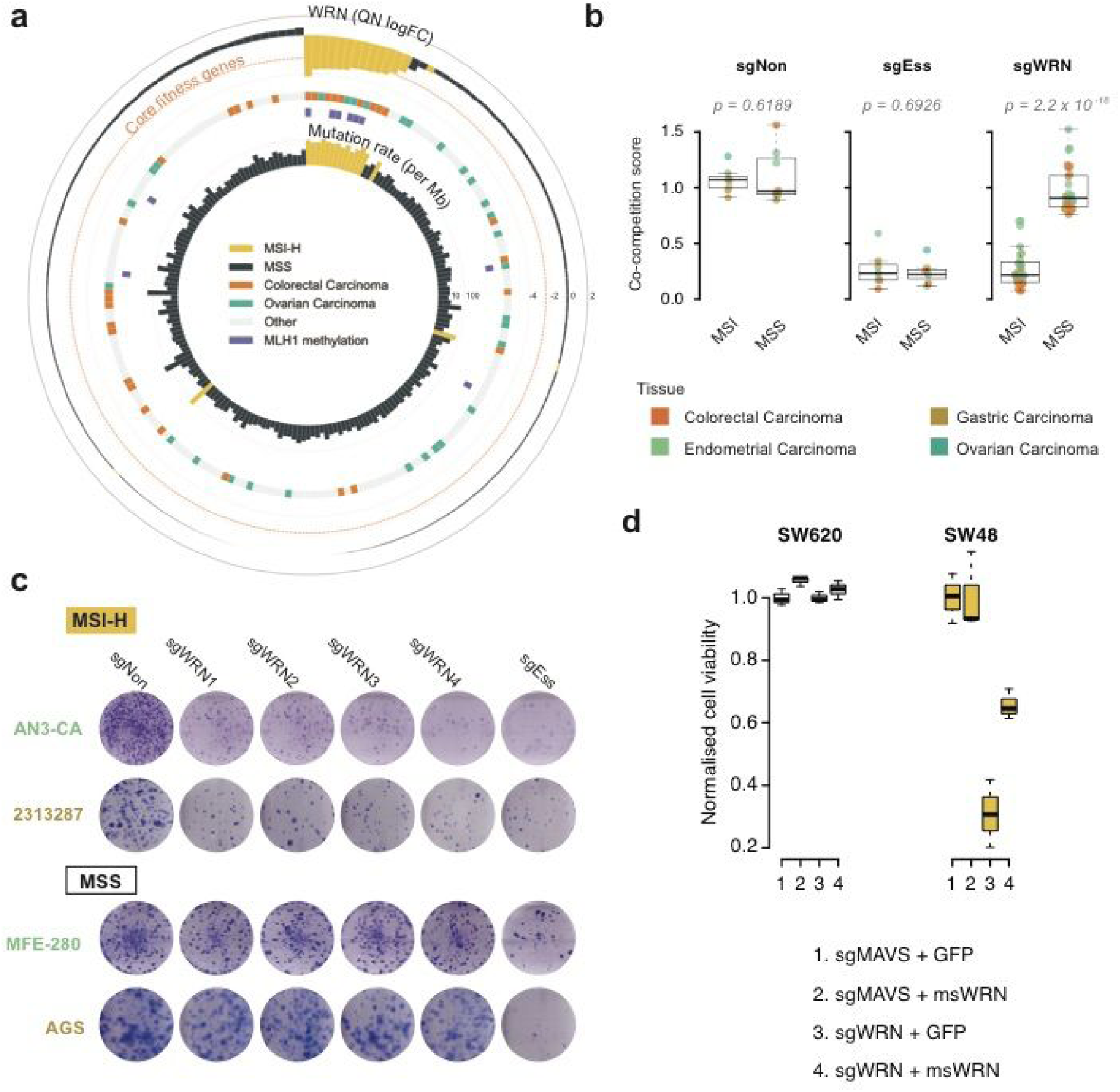
WRN is a target in MSI-H cancer cells from multiple tissue types. (a) Association of *WRN* essentiality with MSI-H, *MLH1* promoter hypermethylation and mutation burden. For comparison the median fitness effect of core fitness genes is shown. (b) Validation of *WRN* dependency using a co-competition assay. sgRNAs targeting an essential (sgEss) and non-essential gene (sgNon) were used as controls. Each point represents the mean co-competition score for a cell line (7 MSI-H and 7 MSS lines assayed in duplicate). (c) Representative clonogenic assays confirmed selective *WRN* dependency in MSI-H gastric and endometrial cell lines. (d) Expression of mouse *Wrn* cDNA from a transgene rescues the loss of fitness associated with *WRN* knockout in MSI-H SW48 cancer cells. No effect is observed in SW620 MSS cells. Mouse *Wrn* cDNA lacks the human sgRNA targeting sequence. sgRNA for mitochondrial antiviral signaling protein (MAVS), a known non-essential gene, and GFP cDNA are used as controls. Data are the average of 3 independent experiments with standard error. Box and whisker plots are median, interquartile range and 95% confidence intervals.

## Discussion

Oncology drug discovery is notoriously difficult and new approaches are needed to more effectively identify and prioritise therapeutic targets. Here, we performed CRISPR-Cas9 screens in a diverse collection of cancer cells lines and combined this with genomic and tractability data to nominate new oncology targets. Our results should facilitate the unbiased selection of a new and broader portfolio of candidate targets for drug development. Equally, they will assist in deprioritising targets lacking supportive evidence. We recognise that not all targets are suitable for drug development, but even a modest improvement in drug development success rates could have considerable impact on reducing drug development costs and bring patient benefit. Beyond the specific application used here, our CRISPR-Cas9 screening results provide the research community with a rich resource with diverse applications in biology and human genetics.

Despite the comprehensive and systematic approach used here, several limitations of our study should be acknowledged. First, we focused on cell-intrinsic dependencies which impact on fitness, and currently do not consider targets involving the tumour environment. Second, although CRISPR-Cas9 screens have high efficiency and specificity, we cannot exclude false-positive and false-negative results ^33^. Third, modulation of a drug target, for example with a small molecule or antibody, may not replicate the effect of genetic deletion mediated using CRISPR-Cas9, and functional redundancy between gene family members may mask essentialities that could be targeted with drugs. Finally, although *in vitro* cell lines are a valuable tool for drug discovery, they do not recapitulate some aspects of *in vivo* tumour biology and consequently may not always be predictive of patient responses. Thus, confirmatory studies are necessary to evaluate candidate targets.

Amongst the top targets, we identified WRN protein as a promising target in MSI-H tumours for multiple cancer-types, indicating that WRN antagonists could in the future be exploited as a targeted therapy in this histology-agnostic molecular subgroup of patients. The mechanism underpinning WRN dependency in MSI cancers is the subject of ongoing investigation but could be due to an interplay between MMR and the role of *WRN* in DNA replication and repair, or possibly a more direct role for *WRN* in MMR ^34^. Mutation of *WRN* leads to the autosomal recessive disorder Werner Syndrome, which is characterised by premature ageing and a median age of death of 54 years ^35^. Thus, although loss of WRN is compatible with human development, given its role in maintaining genome integrity, targeting WRN could result in collateral damage to normal cells and so careful consideration should be given to maximising the therapeutic index through patient selection and dose scheduling. Immune checkpoint inhibitors are approved for the treatment of MSI tumours ^36^, suggesting that a possible route for clinical development of WRN inhibitors would be as an adjunct therapy and in the setting of resistance to immune checkpoint inhibitors.

In summary, we have developed an unbiased and systematic framework that effectively serves to inform ranked priority targets. This framework will further evolve as more screens are performed and additional cancer genomic datasets are integrated. Efforts such as ours to build a compendium of fitness genes in cancer cells, and the identification of context-specific gene dependencies, could transform the decision making process of drug development to improve success rates and ultimately patient outcomes.

## References and Notes

## Acknowledgments

We thank David Adams, George Vassiliou and Leo Parts for their critical reading of the manuscript. FI thanks Ethan Julian Iorio for his insightful comments on the visualisations included in this manuscript.

## Author contributions

Conceptualization: M.G./F.I./K.Y/C.B-D

Methodology: M.G./F.B/K.Y

Software: F.I./E.G.

Validation: G.P./E.G./F.B.

Formal analysis: F.B/F.I/E.G.

Investigation: F.B./G.P./C.B./Y.R/R.A./A.D.J./R.McR./R.P./P.W.

Resources: M.G./F.B./F.I./R.M.S./J.S-R.

Data curation: F.B./F.I./E.G.

Writing – Original Draft Preparation: M.G./F.B./F.I./E.G.

Writing – Review & Editing: M.G./F.B./F.I./E.G./G.P./E.S/K.Y./J.S-R

Visualization:F.B./F.I./E.G.

Supervision: M.G./F.B/C.B./S.H./E.S/D.D./J.S-R./K.Y.

Project administration:F.B./E.S/K.Y./M.G.

Funding Acquisition: M.G./J.S-R./C.B-D/K.Y.

## Data and materials availability

All data is available in the main text or the supplementary materials, software code is available through github (URLs provided in Supplementary Materials).

## Supplementary Materials

Materials and Methods

Table S1 - S13

Fig S1 – S9

