## Supplementary Methods, Supplementary Figures and Legends of Supplementary Tables for "Prioritisation of oncology therapeutic targets using CRISPR-Cas9 screening"

**Affiliations:**

**This PDF file includes:**

Materials and Methods

Figs. S1 to S9

Captions for Tables S1 to S13 (enclosed as Supplementary Data)

### Materials and Methods

#### 1. CRISPR-Cas9 screening

**Plasmids**

All plasmids used have previously been described [1] and are available through Addgene (Cas9 vector- 68343; gRNA vector - 67974). Plasmids were packaged using the ViraPower Lentiviral Expression System (Invitrogen; Cat No. K4975-00) as per manufacturer's instructions.

**Cell culture**

All cell lines used in this study (Table S1) were selected from the Genomics of Drug Sensitivity in Cancer (GDSC):1000 cell line panel [2] and maintained as supplier recommended. To control for and identify cross-contamination and sample swap, a panel of 92 SNPs was profiled for each cell line before and following completion of the CRISPR-Cas9 library screening pipeline.

**Generation of Cas9-expressing cancer cell lines**

Cells were transduced with lentivirus containing Cas9 in T25 or T75 flasks at ~80% confluence in the presence of polybrene (8µg/ml). The cells were incubated overnight followed by replacement of the lentiviral containing medium with fresh complete medium. Blasticidin selection commenced 72 hours post-transduction at an appropriate concentration determined for each cell line using a blasticidin dose response assay (blasticidin range 10-75µg/ml; cell viability assessed using CellTiter-Glo 2.0 Assay (Promega; Cat No. G9241)). Cas9 activity was assessed as described previously [1]. Only cell lines with Cas9 activity >75% progressed to the next stage of the pipeline.

**Genome-wide sgRNA library and screen**

Two genome-wide sgRNA libraries were used in this study: Human CRISPR Library v1.0 and Human CRISPR library v1.1. The Human CRISPR library v1.0 is described previously [1]. Human CRISPR library v1.1 also targets 18,025 genes and contains all sgRNAs from v1.0 plus an additional 1,004 non-targeting sgRNAs. Library v1.1 was synthesised using high-throughput silicon platform technology (Twist Bioscience), resulting in a more uniform representation of individual sgRNAs. A subset of genes in library v1.1 were targeted with an additional 5 sgRNAs. This subset of genes was comprised of kinases, epigenetic related genes and *a priori* known essential genes. Although, for consistency, all the computational analyses were focused on the set of overlapping sgRNAs across the two libraries only, and counts for the additional guides in library v.1.1 where removed at the preprocessing phase and not used in this study. The HT-29 cell line was screened with both libraries and resulting datasets kept separated for comparative analyses (results summarised in in Fig. S2D).

A total of $3.3 \times 10^7$ cells were transduced with an appropriate volume of lentiviral packaged whole-genome sgRNA library to achieve 30% transduction efficiency (100x library coverage). This volume was determined for each individual cell line using a titration of packaged library and assessing the percentage of BFP-positive cells by flow cytometry. Transduction was completed in technical triplicate (or duplicate for cell lines with a large cell size e.g. glioblastoma). Due to the large number of screens performed, multiple batches of packaged library virus were prepared. Each batch was tested in HT-29 to ensure consistency of batch preparations. HT-29 was also screened every 3 months to ensure the quality of data generated by the pipeline was consistent over time. Transduction efficiency was assessed 72 hours post transduction. Samples with transduction efficiency between 15-60% proceeded through the pipeline to puromycin selection. The appropriate concentration of puromycin for each individual cell line was determined from a dose response curve (puromycin range 1-5μg/ml; cell viability assessed using CellTiter-Glo 2.0 Assay (Promega; Cat No. G9241). The percentage BFP-positive cells was reassessed after a minimum of 96 hours of puromycin selection. For samples with <80% BFP-positive cells, puromycin selection was extended for additional 3 days and the percentage BFP-positive cells were assessed again. Cells were maintained until day 14 post transduction with a minimum of $5.0 \times 10^7$ cells reseeded at each passage (500x library coverage). Approximately $2.5 \times 10^7$ cells were harvested, pelleted and stored at -80 °C until genomic DNA extraction.

**DNA extraction, sgRNA PCR amplification, Illumina sequencing and sgRNA counting**
Genomic DNA was extracted from cell pellets using either the QIAsymphony automated extraction platform (Qiagen; QIAsymphony DSP DNA Midi Kit, Cat No. 937255) or by manual extraction (Qiagen; Blood & Cell Culture DNA Maxi Kit, Cat No. 13362) as per manufacturer's instruction. PCR amplification, Illumina sequencing (19-bp single-end sequencing with the custom primer on the HiSeq2000 v4 platform) and sgRNA counting were performed as described previously [1].

### 2. WRN dependency in MSI-H cell lines

**Co-competition Assay**
The sequence of sgRNAs targeting *WRN* and cell lines used in validation experiments are described in Tables S12 and S13, respectively. The sgRNAs were inserted into pKLV2-U6gRNA5(BbsI)-PGKpuro2ABFP-W (Addgene #67974). Cell lines were transduced at ~50% efficiency as described above in 6-well plates. The percentage BFP positive cells was assessed at day 4 and day 14. A co-competition score was determined as the ratio of BFP cells on day 14 compared to day 4. A co-competition score of 1 indicates no reduction in BFP positive cells compared to BFP negative cells (e.g. no fitness effect). A co-competition score below 1 indicates a reduction in cell number following sgRNA transduction.

**Clonogenic Assay**
Cell lines were transduced with lentivirus encoding sgWRN at ~100% efficiency as described above in 6-well plates (2,000 cells/well), typically for 15 - 21 days. Cells were fixed using

100% ice cold ethanol for 30 minutes followed by Giemsa staining overnight at room temperature.

**Western blot analysis**
Cells were transduced at ~100% as described above in 10 cm dishes. Day 5 post transduction, cells were lysed with 200µl RIPA buffer. Lysates were used for SDS-PAGE and immunoblot analysis for WRN was completed using anti-WRN antibody (Cell Signalling Technologies, #4666; dilution 1:2000).

**WRN rescue experiment**
SW620 and SW48 cells (2 x $10^5$ cells) were transfected by nucleofection (Lonza 4D Nucleofector Unit X) with Cas9/sgRNA RNP targeting human *MAVS* gene (used as a non-essential knockout control) or *WRN*, together with overexpression of 200 ng pmGFP control or 200ng mouse *Wrn* cDNA (NM 011721, Origene Cat# MR226496 ). From each sample post nucleofection 5000 cells were seeded in a 96-well and allowed to grow for 5 days, after which cells were collected for either CellTiter-Glo assay (Promega Cat# G9241) or blotted with a WRN antibody (ThermoFisher Cat# PA5-27319). β-actin (Cell Signaling Cat#4970) was used as a loading control in western blot. sgRNA sequences used are listed below:

*WRN* gRNA sequences:
WRN g1: TGGCCACCATTATACAATAG
WRN g2: CATTCATTACGGTGCTCCTA
*MAVS* gRNA sequences:
MAVS g1: CCCTGGCCCGTTCCACCCCC
MAVS g2: AGGCTGGAGGTCGCACCTGC

CellTiter-Glo data was read on a Envision Multiplate Reader and data analysis was performed using GraphPad Prism 7 software. t-test was performed using the multiple t-test module in Prism 7.

### 3. Computational analyses

#### 3.1 CRISPR screen data analyses

**Screening quality control (QC) 1: low-level QC assessment and filtering**
To perform QC assessment and assess reproducibility of screening outputs, correlation scores were computed for each cell line between replicates using sgRNA treatment counts at a genome-wide level (the results shown in Fig. S1C), as a first explorative analysis. To define a reproducibility threshold, we developed an approach based on [3]. Specifically, we restricted the analysis to a set of 505 most informative sgRNAs, identified within sets of sgRNAs targeting the same gene, on the basis of the correlation between corresponding patterns of logFCs across all screened cell lines (R > 0.6). We next computed averaged gene-level

logFCs for 180 genes on the domain of these informative sgRNAs, per each individual technical replicate, and computed all pairwise correlation scores between resulting gene-level profiles of logFCs. This allowed estimating a null distribution of replicate correlations (plotted in gray in Fig. S1D). We then defined a reproducibility threshold as the R value where the estimated probability mass function of the correlation scores computed between replicates of the same cell line was at least twice that of the null mass probability function (Fig. S1D): R = 0.6057. Of the 204 screened cell lines, 187 had an average replicate correlation higher than this threshold, thus passed the reproducibility assessment and for 5 cell lines there were no replicates. Excluding the least reproducible replicate for 5 of the remaining 12 cell lines not passing the first reproducibility assessment allowed their average replicate correlation to exceed the threshold defined above, thus resulting in a final analysis set of 197 cell lines (listed, together with all QC scores in Table S1).

**Screening QC 2: screening performance assessment**

We next considered the genome-wide profiles of gene-level logFCs (averaged across targeting sgRNAs and replicates) of each screened cell line as a classifier of predefined sets of essential/non-essential genes (from [4]), respectively *E* and *N*, by means of Receiver Operating Characteristic (ROC) indicators. The analysis was restricted to genes only belonging to either *E* or *N*, and hereby defined as set *G*. For each cell line we assembled a *predictor* vector *P* containing the logFCs for all the genes in *G*, and a boolean *response* vector with one entry per gene in *G*, which was equal to *TRUE* if the corresponding gene belonged to *E*. Specificity, Precision and Sensitivity curves were computed matching these two vectors, by making use of the *roc* function of the pROC R package [5]. The resulting ROC curves were shown in Fig. 1C and Fig. S1G.

As final QC assessment, we measured the depletion signal magnitude observed in each screened cell line by evaluating the median logFC, and the discriminative distance between their distributions, for predefined essential/non-essential genes from [4], and ribosomal protein genes from [6]. In particular, we computed for prior known essential (*E*) and ribosomal protein (*R*) genes a Glass Δ defined as:

$$| \mu(\text{logFC}(x \in X)) - \mu(\text{logFC}(n \in N)) | / \sigma(\text{logFC}(x \in X)),$$

Where $X \in \{E, R\}$, and $\mu, \sigma$ indicate mean and standard deviation, respectively. Results across all screened cell lines are shown in Supplementary Fig. 1D.

**sgRNA count preprocessing and CRISPR-bias correction**

Samples that passed QC assessment (197 cell lines) were further processed as described below to identify fitness genes. We made use of our *CRISPRcleanR* R package [7], publicly available and fully documented at https://github.com/francescojm/CRISPRcleanR.

sgRNAs with less than 30 reads in the plasmid and sgRNAs belonging to library v1.1 only were removed from further analysis. Remaining sgRNAs were assembled into one file per cell line, including the read counts from the matching library plasmid and all replicates, followed by normalisation using a median-ratio method to adjust for the effect of library sizes and read count distributions [8]. Depletion/enrichment logFCs for individual sgRNAs were quantified between post library-transduction read-counts and library plasmid read-counts at the individual replicate level. This was performed using the *ccr.NormfoldChanges* function of CRISPRcleanR.

Next we performed CRISPR bias correction. For each cell line, sgRNA-level logFCs were averaged across replicates and sorted by genomic coordinates (with the *ccr.logFCs2chromPos* function). We applied a circular binary segmentation algorithm [9,10] to the genome-sorted patterns of sgRNAs' logFCs. This allowed us to identify genomic regions containing clusters of sgRNAs targeting a large number of different genes, with sufficiently equal logFCs, which on average are significantly different from the background. The logFCs of the sgRNAs targeting these regions were mean centered. This was performed using the the *ccr.GWclean* function of CRISPRcleanR with default values for all arguments. Effectiveness of this correction method in reducing false positives (non expressed amplified genes) while keeping constant true positive rates, and distributions of logFCs for *a priori* known essential and non-essential genes is shown in [7]).

**Fitness gene calling**

The CRISPRcleanR-processed profiles of sgRNAs-level logFCs (corrected logFCs) were used as input to an in-house R implementation of the BAGEL method [4] to call significantly depleted genes (code publicly available at https://github.com/francescojm/BAGELR). With respect to the original BAGEL python package, our implementation computes gene-level Bayesian factors (BFs) by averaging those of corresponding targeting sgRNAs, instead of summing them. Additionally, it uses reference sets of predefined essential and non-essential genes described in [4]. However, in order to avoid their status (essential/non-essential) being defined *a priori*, we removed any high-confidence cancer driver genes as defined in [2]. The resulting curated reference gene sets are available as built-in data objects in the R implementation of BAGEL.

A statistical significance threshold for gene-level BFs was determined for each cell line as in [11]: after ranking all the screened genes based on their BFs in decreasing order, from most to least depleted for each rank position $r$, a corresponding set of genes $G_r$ was defined by pooling together all the genes whose rank position was $\leq r$. Then a false discovery rate FDR(

$r$) was computed as $100 \times |G_r \cap N| / |G_r \cap (E \cup N)|$, where $E$ and $N$ are the reference sets of *a priori* known essential and non-essential genes described above, respectively. Finally, the BF of the gene in the rank position $r^* = \max_r \{\text{FDR}(r) < 5\%\}$, was defined as the significance threshold for the cell line under consideration and all the genes with a BF above this threshold were deemed as significantly essential (at a 5% FDR).

Finally in each cell line, each gene was assigned a scaled BF computed by subtracting the BF at the 5% FDR threshold defined for each cell line from the original BF, and a binary fitness score equal to 1 if the resulting scaled BF was > 0.

In addition, bias-corrected sgRNAs treatment counts were derived from the corrected sgRNA=level logFCs (using the *ccr.correctCounts* function of CRISPRcleanR) and used as input to MAGeCK [12] for computing depletion significance via mean-variance modeling. This was performed using the MAGeCK python package (version 0.5.3), specifying in the command line call that no normalisation was required.

At the end of this stage, the following gene-level depletion score matrices were produced in each cell line: raw logFCs, bias-corrected logFCs, BFs, scaled BFs, binary fitness scores and MAGeCK depletion FDRs.

### 3.2 High-level CRISPR screen data analyses

**Pan-cancer and cancer-type core fitness genes using the adaptive Daisy model (ADaM)**
We designed the adaptive Daisy model (ADaM), an heuristic algorithm for the identification of core fitness (CF) genes, implemented it in an R package and made it publicly available at https://github.com/francescojm/ADaM. ADaM is based on the simpler Daisy Model described in [11]. To identify genes that are consistently depleted across multiple cell lines (hence considered as a proxy set of CF genes) the Daisy Model computes a *fuzzy* intersection $I_{m^*}$ (the core of the daisy) composed of genes that are significantly depleted (fitness genes) in at least $m^*$ cell lines, where $m^*$ is apriorististically defined. The genes not belonging to $I_{m^*}$ (thus falling on the petals of the daisy) are deemed as context-specific essential (CSE) genes. In [11] the authors screened 7 cell lines from different tissues and defined genes significantly essential in at least $m^* = 3$ cell lines as CF genes. Generalising this approach, ADaM (i) applies the Daisy Model but adaptively determines $m^*$ in a data-driven way, predicting sets of CF genes (one set per input group), (ii) applies the Daisy Model to the obtained sets of cancer-type CF genes by adaptively defining the number of cancer types $k$ for which a gene should have been predicted as CF in order to be considered as a pan-cancer CF gene. The whole process is illustrated in Fig. S11 for an example cancer type (ovary) and for determining pan-cancer CF genes.

Briefly, for a given cancer type $T$ for which $M$ cell lines have been screened, ADaM computes fuzzy intersections of genes $I_m$, for each $m = 1, \ldots, M$, including genes that are significantly depleted in at least $m$ cell lines. Subsequently for each $m = 1, \ldots, M$, a true positive rate (TPR($m$)) is computed by considering as true positives the genes included in an priori known essential gene set $E$:

$$\text{TPR}(m) = |E \cap I_m| / |E \cap G|,$$

where $G$ is the set of all screened genes.

At the same, time the deviance of $|I_m|$ from its expectation $\pi_m$ is computed as follows:

$$\text{D}(m) = \log_{10}(|I_m| \;/\; \pi_m).$$

To estimate $\pi_m$, 1,000 randomised versions of the binary depletion scores (computed as detailed in the previous section) of all the genes across the cell lines from $T$, are generated while preserving the total number of depleted genes per cell lines, to account for their overall vulnerability. Then $\pi_m$ is defined as the average value of the $|I_m|_i$ (for $i$ = 1, …, 1000) computed across the randomised versions of the binary depletion scores.

Finally, $m^*$ is defined as the maximal value of $m$ providing the best trade-off between TPR($m$) (inversely proportional to $m$) and D($m$) (proportional to $m$, if the distribution of the number of genes depleted in a fixed number of cell lines is bimodal (example in Fig S11D).

We applied ADaM to the depletion scores observed in cell lines from each of the 12 screened cancer types in turn, defining an ADaM threshold $m^*$ for each of them, and considering the corresponding $I_{m^*}$ sets as proxies of cancer-type CF essential genes and those not belonging to $I_{m^*}$ as proxies of context-specific genes.

Finally, we applied the same method to determine a minimal number $k$ of cancer types for which a gene should be predicted as CF genes in order to be considered as a pan-cancer CF gene.

**Characterisation of ADaM pan-cancer CF genes**

Reference sets of essential ($E$) and non-essential ($N$) genes were extracted from [4]. Other reference gene sets (used while characterising the ADaM pan-cancer CF genes, detailed below) were derived from the Molecular Signature Database (MSigDB [13]) and post-processed as detailed in [7]. A more recent set of *a priori* known essential gene was derived from [14]. $E$, $N$ and the MsigDB sets were also used to filter out genes from the selection of those to be included in the analyses of variance and to assign null target priority scores (both detailed below).

Enrichments of $E$ and the MSigDB signatures in the pan-cancer CF genes identified by ADaM were determined via a hypergeometric test. The pan-cancer CF genes not belonging to any of the aforementioned gene sets were further tested for gene family enrichments by: deriving gene annotations using the BioMart R package [15]; and biological pathway enrichments using a comprehensive collection of pathways gene sets from Pathway

Commons [16] (post-processed to reduce redundancies across different sets as detailed in [17]). All enrichment *p* values were corrected with the Benjamini-Hochberg method.

**Comparison between the ADaM pan-cancer core fitness genes and other reference sets of essential genes.**

We compared the set of pan-cancer core fitness genes predicted by ADaM with the BAGEL reference set of essential genes [4], and a more recently proposed larger set of essential genes [14] in terms of size, estimated precision (number of included true positives / number of included genes) and recall (number of included true positives / total number of true positives). In these comparison we used gold-standard essential genes involved in cell essential processes (downloaded from the MSigDB [13] and post-processed as detailed in [7]). In addition, we estimated false discovery rates for the three gene sets (number of included false positives / total number of false positives) considering as false positives genes predicted to be strongly context-specific essential (thus not core-fitness essential) according to an independent publication [18], and using three different confidence levels. In this independent study, the essentiality signal of each gene is analysed across a large dataset obtained from an RNAi screen across hundreds of cancer cell lines, and its tendency to distribute according to a skewed student-t distribution (indicative of that gene being strongly essential in a minority of cell lines) is estimated. In our comparison we considered sets of putative strongly context-specific essential genes (thus false positives) at different level of likelihood of a skewed student-t distribution.

**Basal expression of cancer-type FC genes in normal tissues**

Basal gene median reads per kilobase of transcript per million mapped reads in normal human tissues were download from the GTEx Portal [19], log transformed and quantile normalised on a tissue type basis.

**Analysis of Variance to identify genomic correlates of gene essentiality**

We performed a systematic analysis of variance (ANOVA) to test associations between gene-level logFC of fitness genes and the presence of 381 cancer driver events (CDEs; 105 SNVs and 276 CNVs) [2] or microsatellite instability (MSI) status at pan-cancer as well as individual cancer-type levels. We focused our analysis on the CDEs because these represent a causal link with carcinogenesis and thus increase interpretability of identified associations and facilitate development of genetic biomarkers. ANOVA was performed at pan-cancer or individual cancer-type level. Seven cancer types with at least 10 screened cell lines were analysed. ANOVA was performed using the analytical framework described in [2] and implemented in a Python package [20], publicly available at https://github.com/CancerRxGene/gdsctools.

Each analysis included only genes that were significantly depleted in at least 2 cell lines (from the considered cancer type, or across the whole panel of cell lines for the pan-cancer analysis), but excluding genes in the BAGEL curated $E$ set, the set of ribosomal protein genes and the other MSigDB genes sets (detailed in the previous section). Genes predicted to be pan-cancer or cancer-type CF genes by ADaM were also excluded from respective ANOVA. CRISPRcleanR-corrected gene-level logFC were quantile normalised on an individual cell line basis and used as indicators of gene essentiality.

For each of the genes included in the analysis, an essentiality vector consisting of $n$ depletion logFCs (described above), one entry per cell line (considering the whole panel for pan-cancer analysis, and only cell line from the analysed cancer type otherwise). The model was linear (no interaction terms) with dependant variables represented by the described vector and factors including tissue type (for the pan-cancer analysis only), microsatellite instability status (for the pan-cancer analysis and for those of cancer types with at least 2 positive samples for this feature) and the status of a cancer driver event (CDE, as defined in [2]), one model for each CDE. Differently from the analyses described in [2] here we used a common set of CDEs (the pan-cancer ones) across the different analyses. Only CDEs occurring in at least 3 cell lines were considered and CDEs with identical patterns of positive occurrence were merged together.

For all the tested gene-CDE associations, effect size estimations versus pooled standard deviation (quantified through the Cohen's d), effect sizes versus individual standard deviations (quantified through two different Glass Δs, for the CDE positive and the CDE negative population respectively), CDE $p$ values and all the other statistical scores were obtained from the fitted models. A CDE-gene pair was tested only if at least three cell lines were contained in the two sets resulting from the dichotomy induced by the CDE-status (i.e. at least 3 positive cell lines and at least 3 negative cell lines), for the pan-cancer and all the cancer-type-specific analyses as well.

The resulting $p$ values were corrected (all together those obtained in the pan-cancer analysis and on a cancer type basis those obtained in a given cancer-type-specific analysis), with the Tibshirani-Storey method [21]. A $p$ value threshold of $10^{-3}$ and a false discovery rate threshold equal to 25% were finally used to call significant associations across all the performed analyses.

#### 3.3. Target priority score

Each gene $t$ was assigned a target priority score $P(t)$ defined as:

$$P(t) = 0.3\, L_1(t) + 0.7\, L_2(t)$$

To formally define the two individual terms $L_1(t)$ and $L_2(t)$, we introduce the boolean function $f(p)$ which evaluates the status of the property $p$ related to a given gene and possibly a given cell line. This functions assumes a value equal to 1 when $p$ is true and equal to 0 otherwise. Additionally, let us consider the following properties:

$p_1(t)$ = {$t$ is involved in at least one ANOVA interaction at FDR < 30% with a CDE or with MSI status at FDR < 5%},

$p_2(t)$ = {$t$ is involved in at least one ANOVA association with a CDE with a *p-value* < 0.001},

$p_3(t)$ = {$t$ is involved in at least one ANOVA association with a CDE with a *p-value* < 0.05},

$p_4(t)$ = {$t$ is involved in at least one *t-test* association with a CDE with a *p-value* < 0.05},

where MSI is microsatellite instability. We used different levels of stringency between the associations with CDEs or MSI due to the number of tests performed in the former being 6 orders of magnitude larger than the latter. The ANOVA associations are those specific to the test group under consideration. In the pan-cancer analysis, each of the properties listed above must satisfy the additional constraint that the considered ANOVA/t-test has to result in at least one of the two Glass Δs (quantifying the effect size with respect to the standard deviations of the two involved sub-populations of samples) > 2. This is to account for the significantly larger number of samples analysed in the pan-cancer setting, which might result in largely significant $p$ values even for small effect size associations.

$L_1(t)$ is then defined as follows:

$$L_1(t) = g(t)\,\{0.8 \sum_{i=1}^{4} f(p_i(t)) +$$

$$0.2 f(\,t \text{ is somatically mutated in at least 2\% of } T \text{ matched primary tumours})\},$$

where somatic mutations from large cohorts of primary tumours were derived from [2] and $g(t)$ is a filter function defined as follows:

$g(t)=\{f(t$ is targeted by more than 1 sgRNA in the employed library) ×

$f(t$ does not belong to any reference set of predefined essential genes) ×

$f(t$ is not predicted as pan-cancer CF gene by ADaM) ×

$f(t$ is not predicted as $T$-specific CF gene by ADaM)},

where the reference set of predefined essential genes are described in the previous section and the last factor is omitted for pan-cancer priority scores.

Finally, $L_2(t)$ is defined as follows:

$$L_2(t) = \sum_c h(t,c) / \sum_c f(t \text{ is significantly essential in } c),$$

where $c$ are the screened cell lines from $T$ (or in the whole panel for pan-cancer scores), essentiality is meant at a BAGEL 5% FDR (as detailed in the previous sections). To defined a $h(t,c)$ let us introduce the following properties:

$q_1(t,c)$ = {scaled BF of $t$ in $c > 1$},

$q_2(t,c)$ = {scaled BF of $t$ in $c > 2$},

$q_3(t,c)$ = {scaled BF of $t$ in $c > 3$},

$q_4(t,c)$ = {MAGeCK depletion FDR of $t$ in $c < 10\%$},

$q_5(t,c)$ = {MAGeCK depletion FDR of $t$ in $c < 5\%$},

$q_6(t,c)$ = {$t$ is highly expressed in $c$ at the basal level},

$q_7(t,c)$ = {$t$ is somatically mutated in $c$},

$q_8(t,c)$ = {$t$ belongs to a biological pathway that is statistically enriched among the genes significantly depleted in $c$},

where the scaled BF and the MAGeCK depletion FDRs are indicators of gene essentiality and are computed in the previous section (Fitness gene calling), $t$ falls over the 95% quantile of basal expression in $c$ according to the FPKM values derived from [22] as described in the following section, somatic mutations for all the cell lines are derived from [2], and pathway enrichments in the set of genes significantly depleted in $c$ are computed with a hypergeometric test using pathways gene sets from Pathway Commons [16] post-processed to reduce redundancies across different sets as detailed in [17].

$h(t,c)$ is then defined as follows:

$$h(t,c) = l(t,c)\,\{0.125 \sum_{i=1}^{8} f(q_i(t,c))\},$$

where $h(t,c)$ is a filter function defined as follows:

$h(t,c) = \{f(t$ is significantly essential in $c) \times$

$f(t$ is express in $c) \times$

$f(t$ is not homozygously copy number deleted in $c)\}$,

where, as before, significant essentiality is meant at a BAGEL 5% FDR, a gene is meant not expressed in a cell line if its FPKM is < 0.05 in that cell line, the gene level copy number status across screened cell lines has been downloaded from (www.cancerrxgene.org, [23]).

#### 3.4 Genome-wide assessment of Target Tractability

To estimate the likelihood of a target to bind a small molecule (SM), or the likelihood of a target to be accessible to an antibody, we have made use of the genome-wide target tractability assessment pipelines developed at GSK. As described in Brown et al [24] the high throughput *in silico* pipeline integrates data from public sources, and assigns human protein-coding genes into hierarchical qualitative buckets. Predicted tractability and confidence in the data increase from Bucket 10 to Bucket 1, with targets in Bucket 1 being considered the most tractable. Of note, targets in lower Buckets (i.e. Buckets 10 to 8) are considered to have uncertain tractability, and should not be ruled out as "intractable" without a deep tractability assessment. This was assembled using data from the following resources.

For the SM pipeline:

- Uniprot [25], PDBe [26], InterPro [27], Pfam [28] & GO [29]: accessed on 18/11/2016
- Complex portal [30], Biomodels [31] accessed on: 24/06/2017
- ChEMBL [32]: version 22 (October 2016)
- SureChEMBL [33]: SureChEMBL RDF as available in Open PHACTS (version 1.2, March 2015)
- Human genome: NCBI [34], November 2016 (20 912 protein coding genes)

For the antibody pipeline:

- HPA [35]: accessed on February 2017
- Uniprot & GO: accessed on 13/07/2017
- Complex portal & Biomodels: accessed on 24/06/2017
- ENSEMBL Compara [36]: accessed on 13/07/2017 (ENSEMBL 89)
- ChEMBL: version 23 (May 2017)
- SureChEMBL: SureChEMBL RDF as available in Open PHACTS (version 1.2, March 2015)
- Human genome: NCBI, November 2016 (20 912 protein coding genes)

#### 3.5 Other computational analyses

##### Collective test for genomic markers of gene essentiality shared among tissues

All the targets for which in at least two cancer types the following condition was satisfied:

$$\sum_{i=1}^{4} f(p_i(t)) \geq 3,$$

i.e. they were associated with at least one CDE *g* (or with the MSI status) at an ANOVA *p* value < 0.001, had their differential essentiality re-tested against the status of *g* in a collective *t-test* considering all the cell lines from the cancer types in which the condition above held, pooled together, yielding the results shown in Fig. S7B and Table S8.

**Characterisation of target protein families and enrichments**

To characterise protein families and compute statistical enrichments we made use of the Panther online tool [37].

**Gene expression data source for and *GPX4* differential expression analysis**

RNA-Seq voom [38] transformed gene-expression measurements were obtained from [22].

For the *GPX4* case, cell lines were divided into two groups according to their loss-of-fitness response to *GPX4* knockout (using a BAGEL FDR < 5% as significance threshold for gene depletion) and gene expression fold-changes were calculated between the *GPX4* essential and non-essential cell lines. Differential gene-expression was statistically assessed using R package Limma [39]. Gene-set enrichment analysis was performed using ssGSEA [13] and cancer hallmark gene sets were used to identify significant enrichment amongst the top differential expressed genes. 10,000 random permutations were performed for each signature to calculate empirical p-values, and FDR correction was applied.

The code used to perform the analysis is readily available and documented as a GitHub project (https://github.com/EmanuelGoncalves/prioritize_crispr).

### Supplementary Figures

**Fig. S1.**

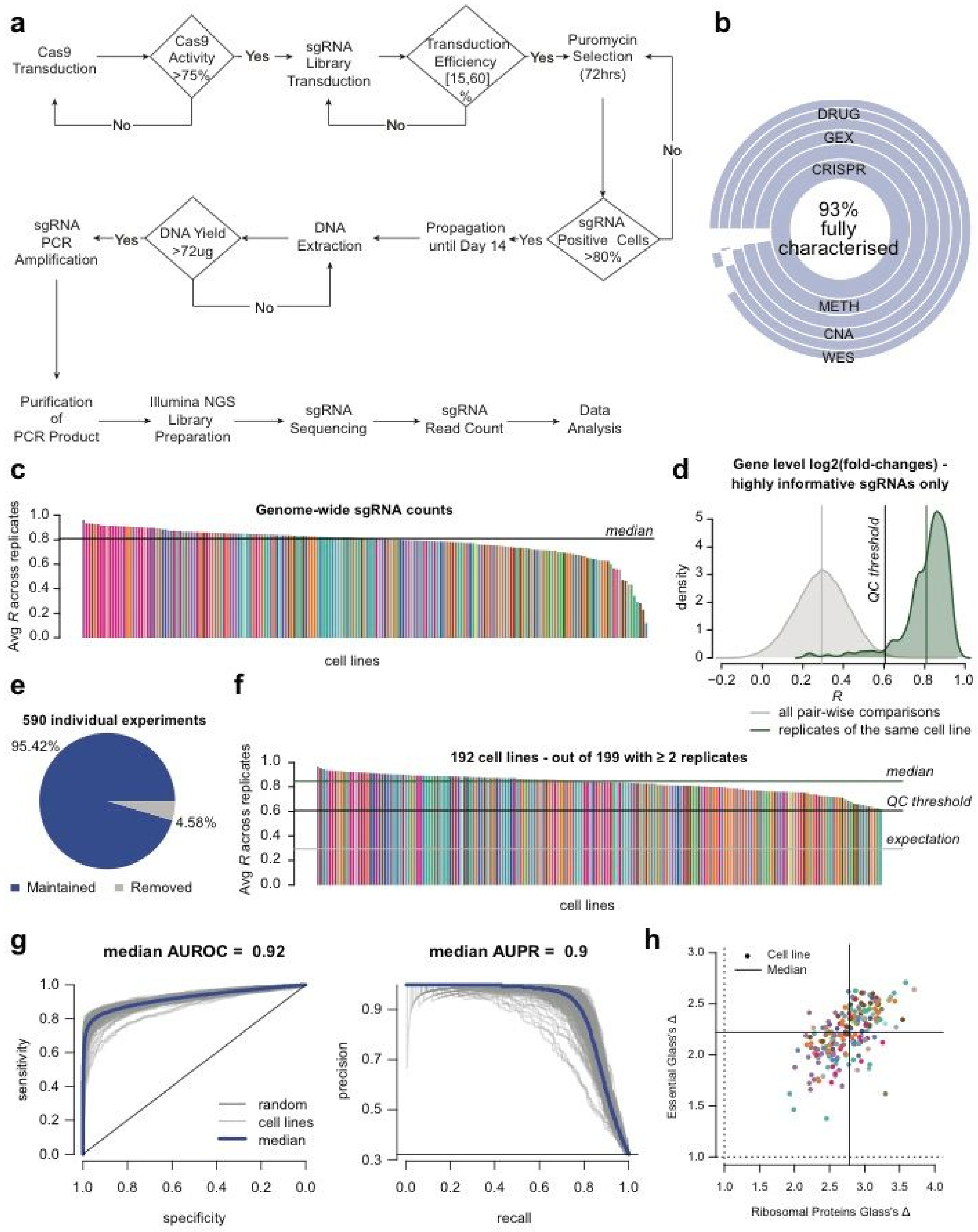

**Fig. S1. CRISPR-Cas9 screening pipeline and data quality.** (a) Schematic of the experimental CRISPR-Cas9 screening pipeline including quality control (QC) checks and go/no-go decisions. (b) Level of genomic characterisation of the CRISPR-Cas9 screened cell lines. (c) Average correlation of replicate sgRNA counts across cell lines. (d) Data QC threshold definition: distributions of correlation scores at the level of sgRNAs' fold-changes computed between replicates of the same cell line (in green) and all possible comparisons (in gray) considering only highly reproducible sgRNAs (detailed in the methods). Vertical lines indicate the mean of the two distributions (in gray and green), and the QC threshold is the R score discriminating between the two distributions. (e) Percentage of cell lines passing/not-passing the data QC filter based on the threshold defined in D. (f) Correlation scores as described in D for cell lines in the final analysis set. (g) ROC and Precision/Recall curves obtained while classifying predefined sets of essential and non-essential genes, based on depletion logFC rank positions; median areas under the curves across all cell lines are reported. (h) Glass delta scores, quantifying the depletion effect size for ribosomal protein genes and *a priori* known essential genes (coordinates on the two axes), across cell lines.

**Fig. S2**

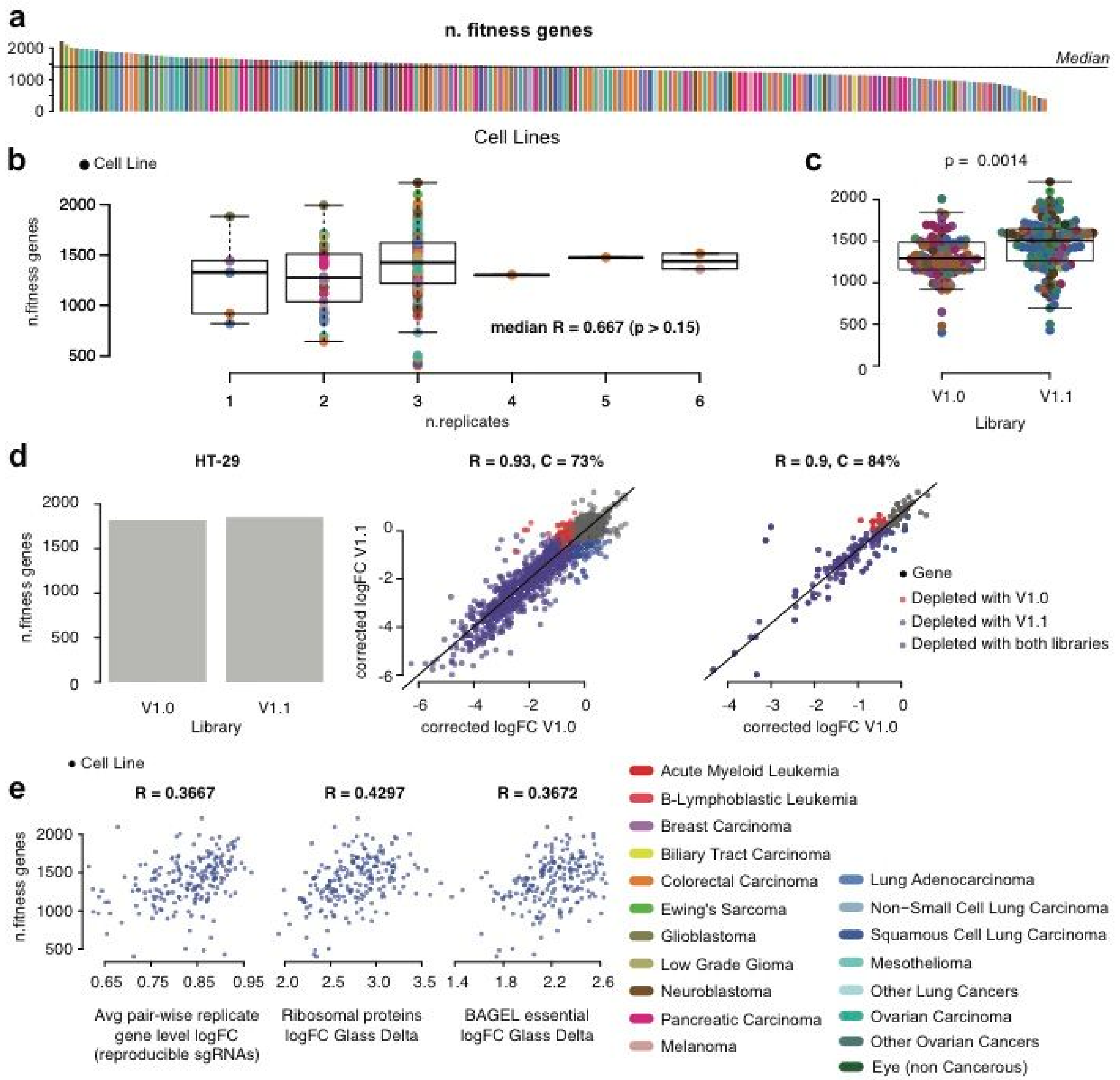


**Fig. S2: Identifying fitness genes in cancer cell lines.** (a) The number of significant fitness genes in each cell line. (b) Correlation between number of screen replicates per cell line (x-axis) and number of significant fitness genes (y-axis). Each dot is a cell line and box-and-whisker plots indicate median, interquartile range and 95 percentiles. A wide variation in the number of essential genes per cell line was observed within each group and a weak non-significant correlation was observed only at the level of median number of fitness genes across groups. (c) The effect of the sgRNA screening library version and the number of fitness genes identified. As expected, an improved version of the library (v.1.1) yields moderately larger numbers of fitness genes, however this is equally variable in both groups and confounded by the tissue of origin of the cell lines. In panels A, B and C, colours indicate cell lines' tissue type. (d) Reproducibility of calling fitness genes in HT-29 cells screening with both sgRNA libraries. Barplot shows number of fitness genes detected and scatter plots depletion scores at genome-wide level or considering only highly informative sgRNAs (as for Fig. S1). In both cases, Fisher Exact test p-value is < $10^{-16}$. (e) Weak correlation between the number of fitness genes per cell line and ability to detect three *a priori* defined sets of essential genes (as detailed in the methods - QC assessment section).

**Fig. S3**

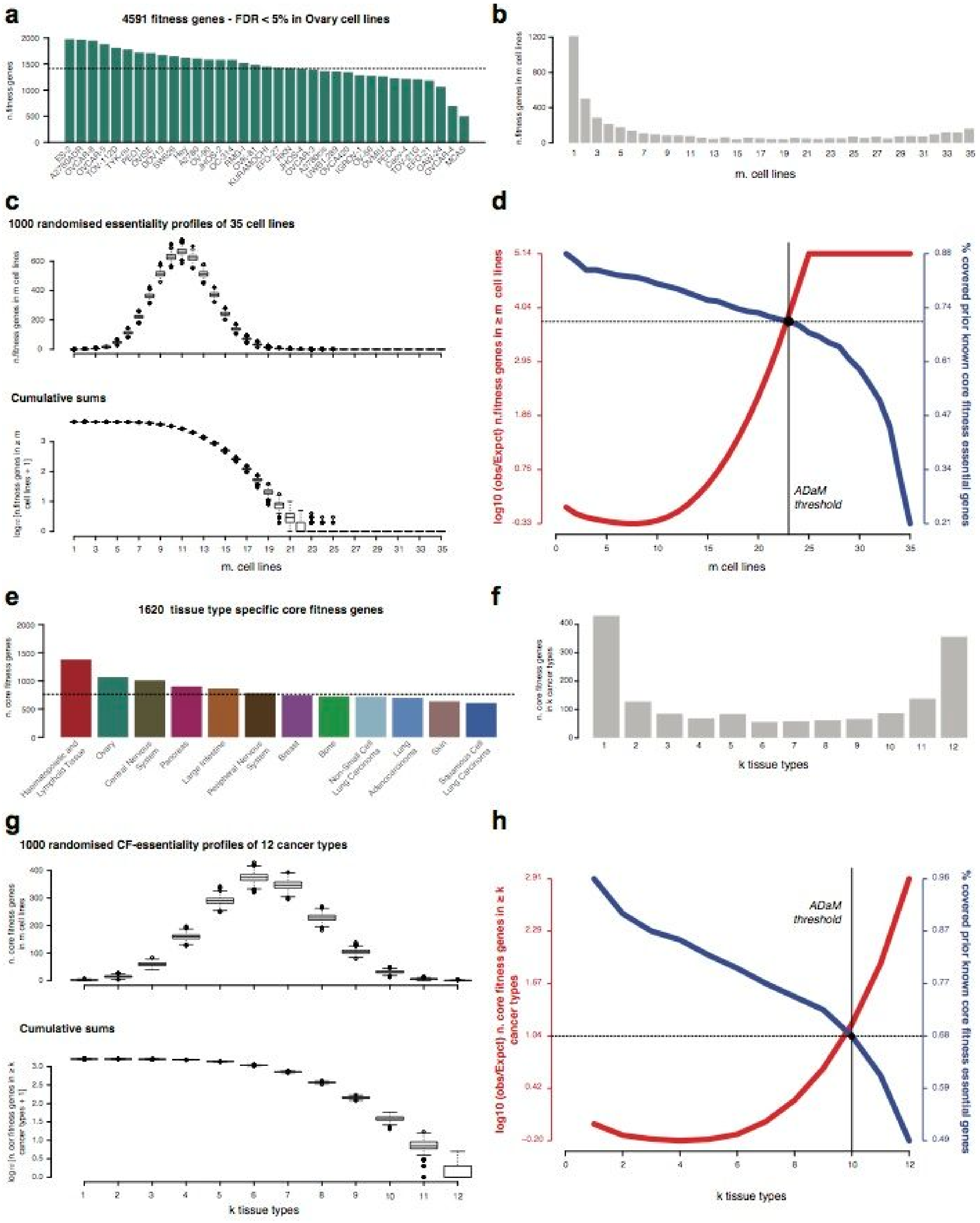

**Fig. S3: Computation of ovary specific (as an example) and pan-cancer core fitness genes with the adaptive daisy model (ADaM).** (a) Number of fitness genes in ovary cell lines. (b) Number of fitness genes observed in a fixed number *m* of cell lines, across numbers of cell lines. (c) Distributions and cumulative distributions of number of fitness genes observed in a fixed number of *m* cell lines, across numbers of cell lines, across 1,000 randomised versions of the depletion scores for ovary cell lines. (d) True positive rates (where positive are *a priori* known essential genes) when considering as predictions the genes that are depleted (fitness genes) in at least *m* cell lines (blue curve), and (red curve) deviance of the number of these gene from expectations (computed as shown in C), across all possible *m* values. The x-coordinate (rounded by excess) of the intersection of these two curves estimates the minimal number of cell lines *m** in which a gene should be significantly depleted in order to be predicted as a core-fitness (CF) gene for the tissue type under consideration. (e) Number of tissue specific CF genes across tissue types. (f) Number of genes predicted as tissue specific CFs for a fixed number *k* of tissue types, across numbers of tissue types. (g) Distributions and cumulative distributions of number of CF genes predicted for a fixed number of *k* tissue types, across numbers of tissue types, across 1,000 randomised versions of the tissue specific CF profiles. (h) True positive rates (where positive are *a priori* known essential genes) when considering as predictions the genes that CFE for at least *k* tissue types (blue curve), and (red curve) deviance of the number of these gene from expectations (computed as shown in g), across all possible *k* values. The x-coordinate (rounded by excess) of the intersection of these two curves estimates the minimal number of tissue types *k** for which gene should have been predicted as tissue specific CF in order to be predicted as pan-cancer CF gene.

Fig. S4

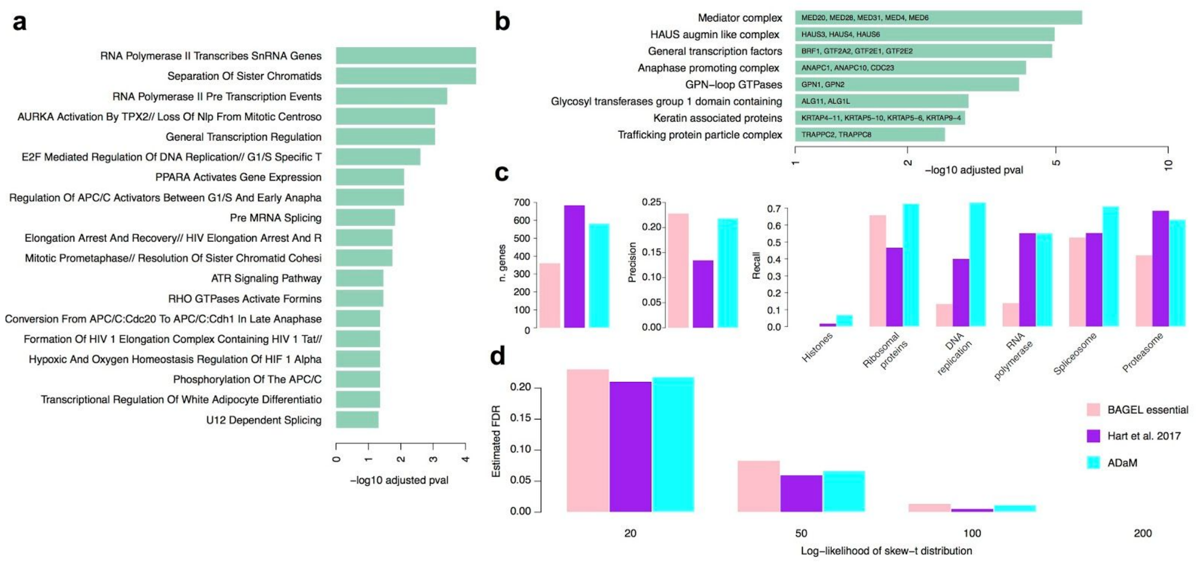


**Fig. S4: Characterization of core fitness genes.** (a) Pathways and gene families (b) statistically enriched (adjusted p-value $< 0.05$) in the core fitness genes newly identified by ADaM. (c) Results from a comparison of the ADaM core-fitness (CF) genes, and two previously reported reference sets of essential genes, in terms of number of genes, estimated precision and recall (considering as true positive the genes included in reference gene sets corresponding to cellular essential process). (d) False discovery rates of putative context-specific essential genes (from an independent study, at different thresholds of reliability) for the ADaM CF genes and two previously reported reference sets of essential genes.

Fig. S5

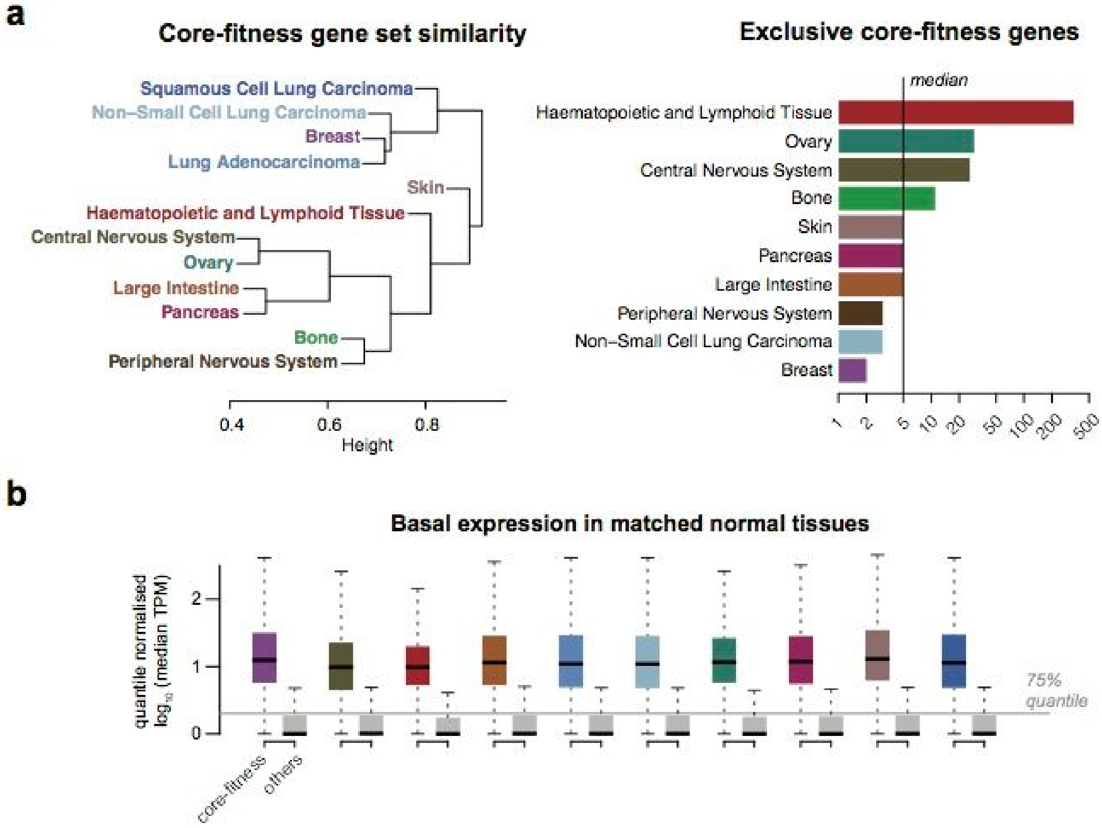


**Fig. S5:** (a) Clustering of tissues based on core fitness gene similarity (left) and the numbers of tissue-exclusive core fitness genes (right). (b) Basal expression of tissue-specific core fitness genes in matched normal tissues is higher than other genes.

**Fig. S6**

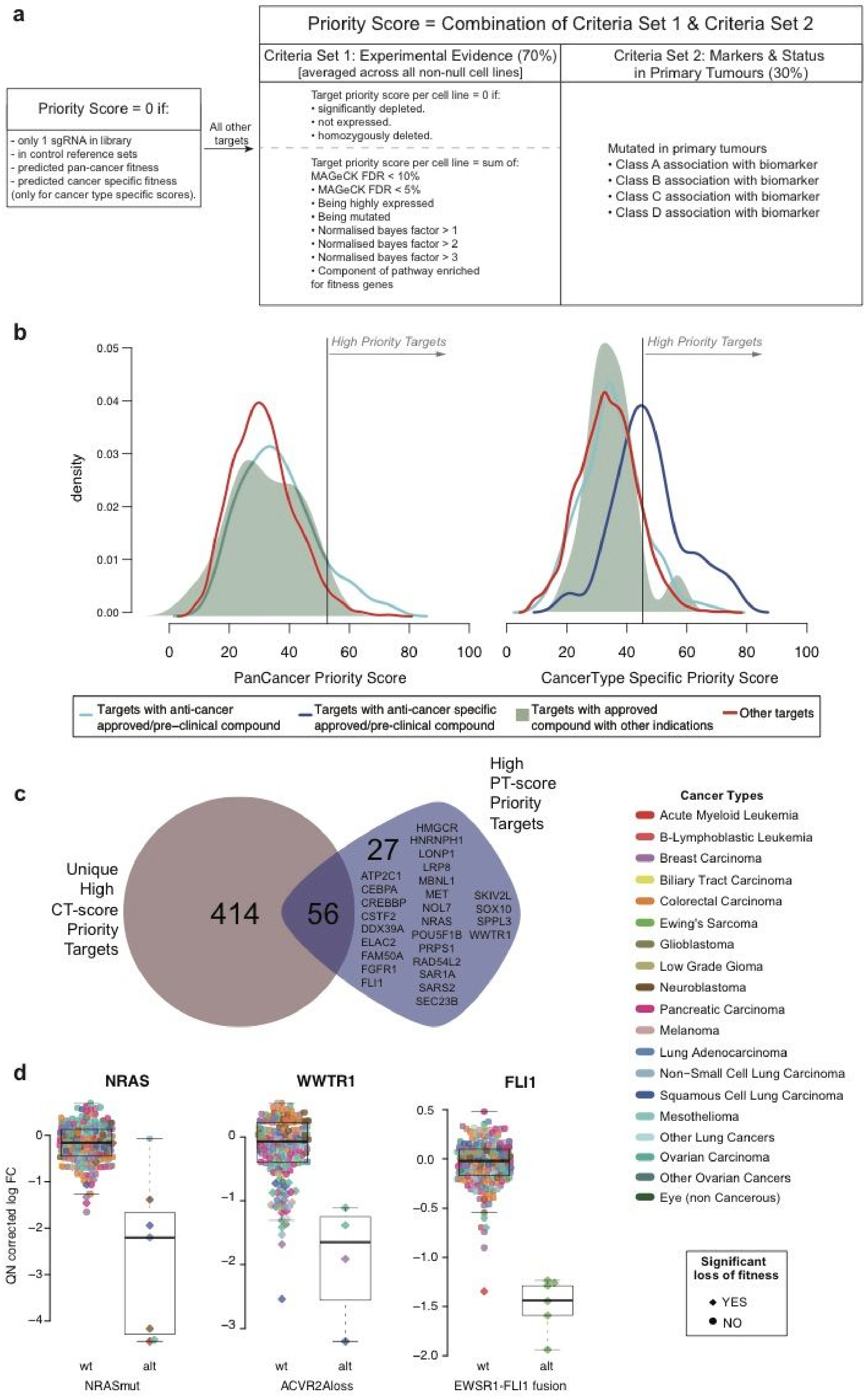
a
Priority Score = Combination of Criteria Set 1 & Criteria Set 2
Criteria Set 1: Experimental Evidence (70%) [averaged across all non-null cell lines]
Criteria Set 2: Markers & Status in Primary Tumours (30%)
Priority Score = 0 if:
- only 1 sgRNA in library
- in control reference sets
- predicted pan-cancer fitness
- predicted cancer specific fitness (only for cancer type specific scores).
All other targets
Target priority score per cell line = 0 if:
• significantly depleted.
• not expressed.
• homozygously deleted.
Target priority score per cell line = sum of:
MAGeCK FDR < 10%
• MAGeCK FDR < 5%
• Being highly expressed
• Being mutated
• Normalised bayes factor > 1
• Normalised bayes factor > 2
• Normalised bayes factor > 3
• Component of pathway enriched for fitness genes
Mutated in primary tumours
• Class A association with biomarker
• Class B association with biomarker
• Class C association with biomarker
• Class D association with biomarker
b
High Priority Targets
density
PanCancer Priority Score
CancerType Specific Priority Score
Targets with anti-cancer approved/pre-clinical compound
Targets with anti-cancer specific approved/pre-clinical compound
Targets with approved compound with other indications
Other targets
c
Unique High CT-score Priority Targets
High PT-score Priority Targets
414
56
27
HMGCR HNRNPH1 LONP1 LRP8 MBNL1 MET NOL7 NRAS POU5F1B PRPS1 RAD54L2 SAR1A SARS2 SEC23B
ATP2C1 CEBPA CREBBP CSTF2 DDX39A ELAC2 FAM50A FGFR1 FLI1
SKIV2L SOX10 SPPL3 WWTR1
Cancer Types
Acute Myeloid Leukemia
B-Lymphoblastic Leukemia
Breast Carcinoma
Biliary Tract Carcinoma
Colorectal Carcinoma
Ewing's Sarcoma
Glioblastoma
Low Grade Gioma
Neuroblastoma
Pancreatic Carcinoma
Melanoma
Lung Adenocarcinoma
Non−Small Cell Lung Carcinoma
Squamous Cell Lung Carcinoma
Mesothelioma
Other Lung Cancers
Ovarian Carcinoma
Other Ovarian Cancers
Eye (non Cancerous)
d
NRAS
WWTR1
FLI1
QN corrected log FC
wt
alt
NRASmut
ACVR2Aloss
EWSR1-FLI1 fusion
Significant loss of fitness
YES
NO

**Fig. S6: Pan-cancer and cancer-type specific priority scores** (a) Details of criteria for the target prioritization scoring system. (b) Distributions of pan-cancer (left) and cancer-specific (right) non-null target priority scores based on the therapeutic indication of approved/pre-clinical compound. A threshold for significance was based on the distribution of scores for targets with approved anti-cancer compounds (specific anti-cancer compounds for the cancer type specific priority score) versus scores for targets with no available compounds. (c) Overlap between cancer-type specific high priority targets (for at least one cancer type) and pan-cancer high priority targets. (d) Example of targets that are scored as high priority only in the pan-cancer context. Each symbol is an individual cell lines colored by tissue type and symbol shapes indicates a significant vulnerability in the cell line.

Fig. S7

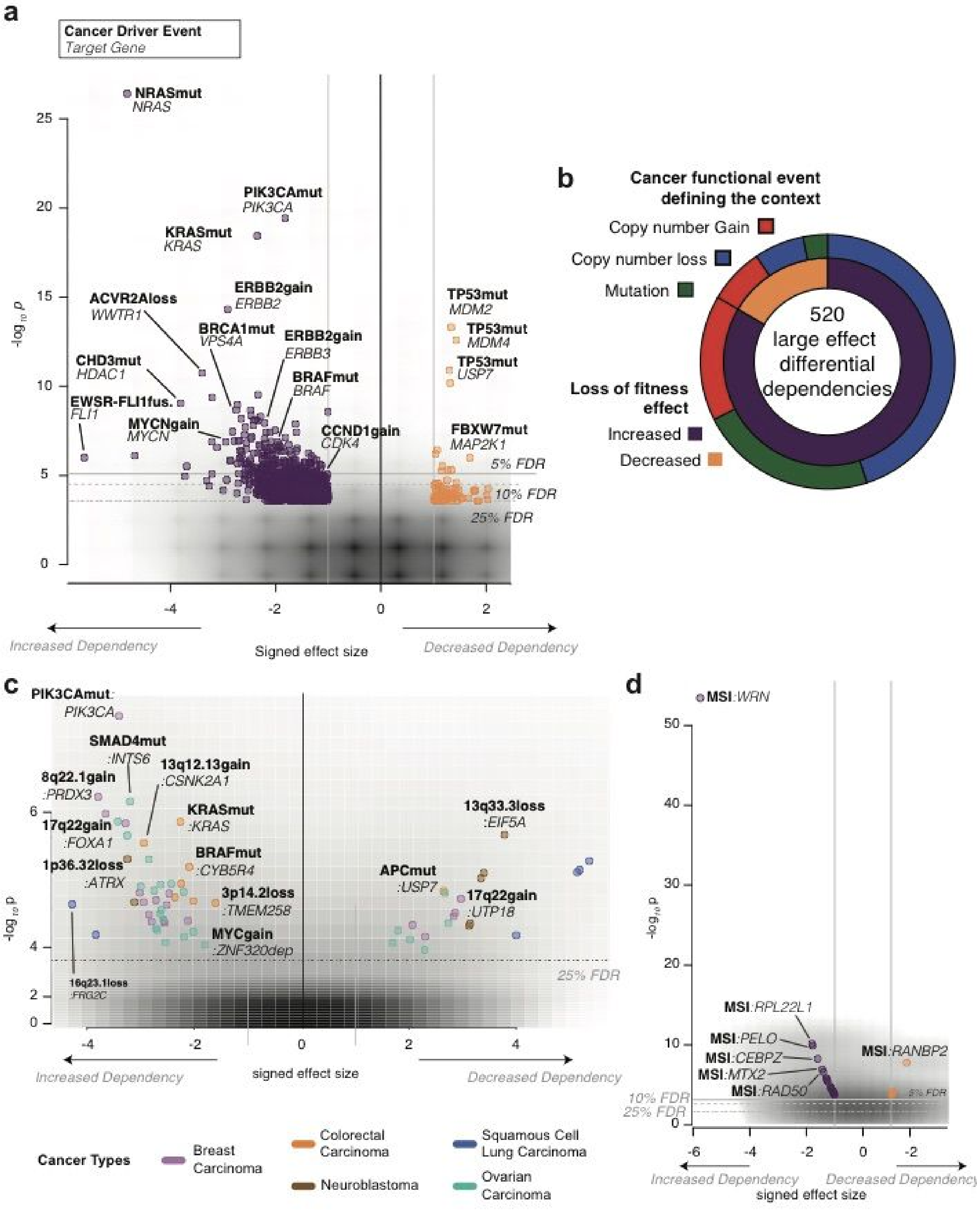


**Fig. S7: ANOVA analysis identifies context specific fitness effects.** (a) Volcano plot with pan-cancer ANOVA results showing statistically significant associations between cancer driver events (CDE) and differential dependency on fitness genes. Each circle represents an association plotted against the signed ANOVA effect size (Cohen's D) (x-axis) and statistical significance (y-axis). CDEs associated with increased or decreased dependency are colored purple and orange, respectively. (b) Summary of large-effect differential dependencies identified through a pan-cancer ANOVAs. The inner circle indicates whether the association is for increased or decreased fitness and the outer circle the class of CDE involved. (c) Results of cancer-type specific ANOVAs between cellular fitness effects and cancer driver events (CDE).  with eEach circle is colored by tissue type and represents an individual association plotted against the signed effect size (x-axis) and statistical significance (y-axis). CDE include recurrent regions of copy number gain or loss and driver genes within each region are described.  (d) Associations between microsatellite instability status of cell lines and gene fitness. (C) Representative examples of cancer-type specific differential dependency on fitness genes when stratified by CDE. Each symbol represents the fitness effect (logFC) in an individual cell line. Symbols are colored by tissue type and shape indicates whether a fitness gene induces a significant vulnerability in the cell line. Box-and-whisker plots indicate median, interquartile range and 95 percentiles. (bD) Summary of large-effect differential dependencies. The inner circle indicates whether the association is for increased or decreased fitness and the outer circle the class of CDE involved.

Fig. S8

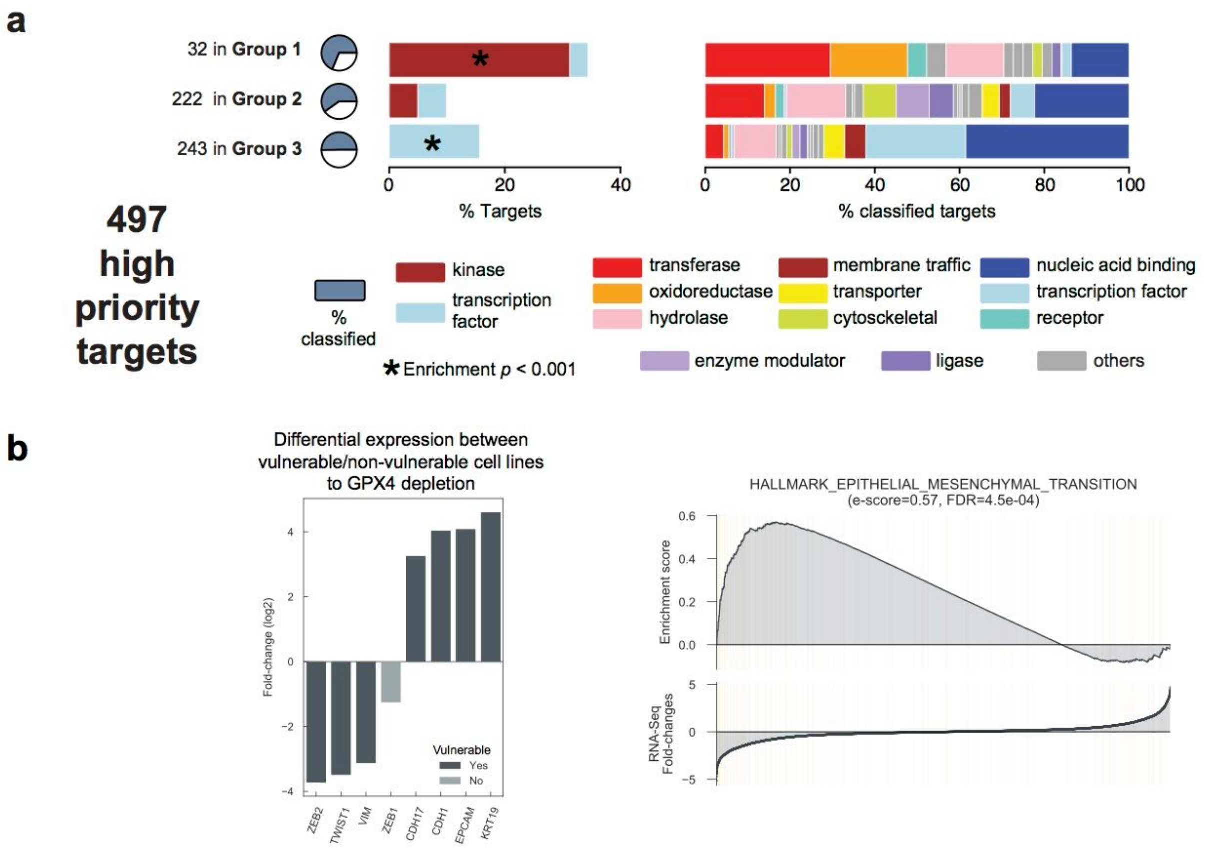


**Fig. S8: Functional classification of priority targets, and GPX4 fitness selectivity for cells undergoing EMT.** (a) Functional classification of priority targets by tractability group. For clarity, kinases (a subset of transferases) and transcription factors are shown separately. Protein classes are indicated by color. Statistical enrichment was calculated using a hypergeometric test. Pie charts on the left indicate the percentage of targets classified in protein families. (b) Genes differentially expressed in cell lines that are vulnerable to GPX4 depletion versus others (left), and top differentially enriched gene signature in cell lines vulnerable to GPX4 depletion.

**Fig. S9**

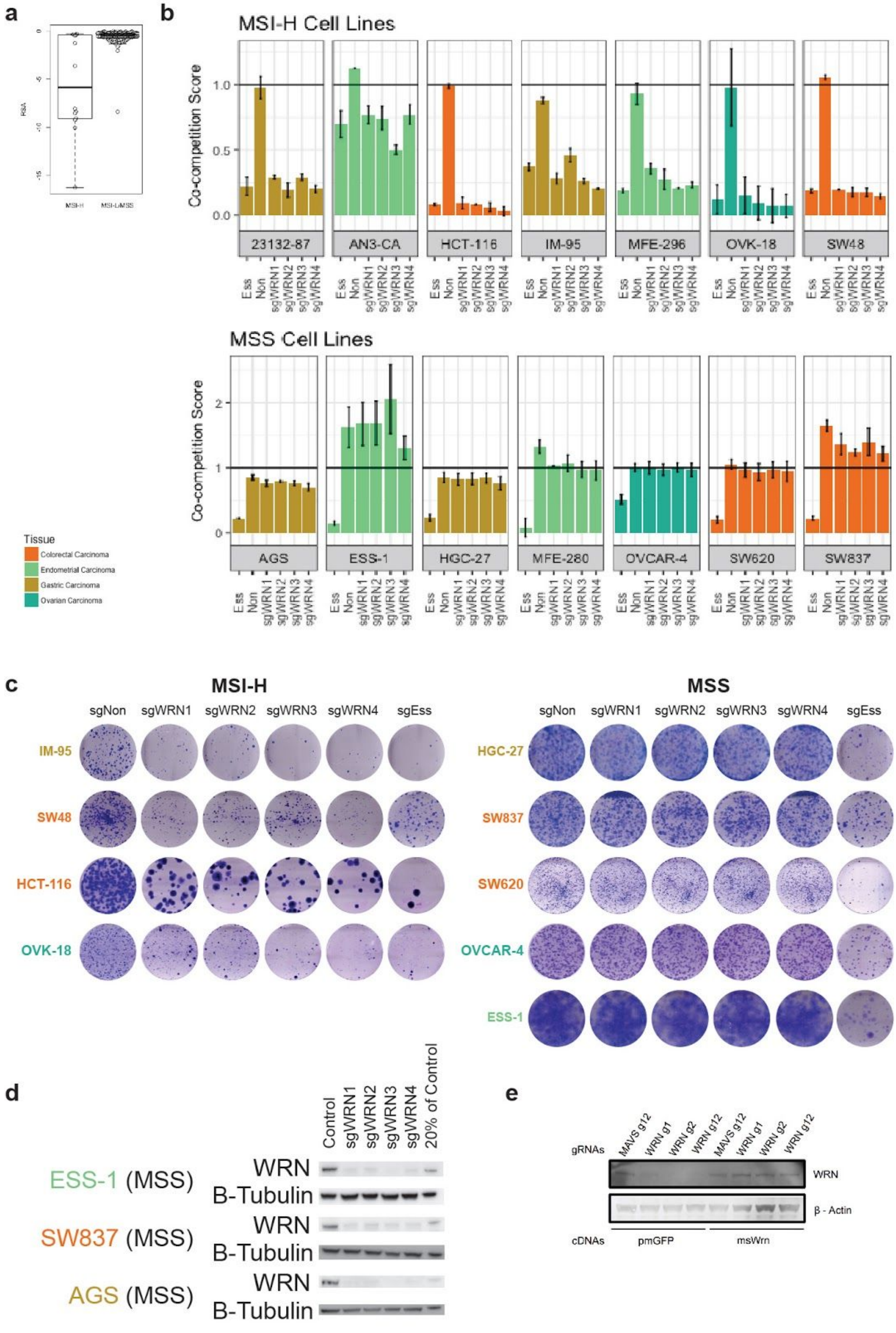
a
b
MSI-H Cell Lines
Co-competition Score
0.0
0.5
1.0
23132-87
AN3-CA
HCT-116
IM-95
MFE-296
OVK-18
SW48
Ess
Non
sgWRN1
sgWRN2
sgWRN3
sgWRN4
MSS Cell Lines
0
1
2
AGS
ESS-1
HGC-27
MFE-280
OVCAR-4
SW620
SW837
Tissue
Colorectal Carcinoma
Endometrial Carcinoma
Gastric Carcinoma
Ovarian Carcinoma
c
MSI-H
sgNon
sgWRN1
sgWRN2
sgWRN3
sgWRN4
sgEss
IM-95
SW48
HCT-116
OVK-18
MSS
HGC-27
SW837
SW620
OVCAR-4
ESS-1
d
Control
sgWRN1
sgWRN2
sgWRN3
sgWRN4
20% of Control
ESS-1 (MSS)
SW837 (MSS)
AGS (MSS)
WRN
B-Tubulin
e
gRNAs
MAVS g12
WRN g1
WRN g2
WRN g12
WRN
β - Actin
cDNAs
pmGFP
msWrn

**Fig. S9: Validation of WRN as a target in MSI-H cancers** (a) The association between *WRN* dependency and MSI status was confirmed using data from an independent RNAi study, Project DRIVE (p-value = 0.004). Each circle represents the RNAi dependency score in an individual cell line. (b) Four sgRNAs targeting *WRN* demonstrated a consistent loss of fitness effect in MSI-H cell lines (7 cell lines; 4 tissue types). This fitness effect was never observed in MSS cell lines (7 cell lines; 4 tissue types). sgRNA targeting non-essential and known essential genes were used as controls. (c) The association between *WRN* loss of fitness and MSI-H status was confirmed using a clonogenic assay in 4 tissue types. (d) A reduction in WRN protein expression following treatment with all sgRNAs targeting WRN was confirmed using Western blot. (e) Western blot confirmation of *WRN* knockout and mouse Wrn transgene expression in MSS cells.

**Captions of Supplementary Tables**

**Table S1:** Cell line annotation, level of characterisation, CRISPR-Cas9 screening details, quality control assessment scores and final analysis set specification.

**Table S2:** Gene fitness summary: binary scores for 6,830 fitness genes across all cell lines.

**Table S3:** Detailed annotation of the essentiality of 6,830 fitness genes including memberships to predefined sets of core fitness essential genes, output of the Adaptive Daisy Model (status of each gene in each cancer type: core fitness, context specific, non-essential) and percentages of vulnerable cell lines (pooled together or within each cancer type).

**Table S4:** Characterisation of pan-cancer core fitness genes. (A) Gene families area under the ROC curves (and significance) using as classifier the rank position of all the genes based on the their n. of vulnerable cell lines, regardless the output of the Adaptive Daisy Model (ADaM). (B) Enrichments of known essential genes used as input to ADaM across the pan-cancer core-fitness genes predicted by ADaM. (C) Full set of pan-cancer core-fitness genes predicted by ADaM with annotation and memberships to the gene sets listed in B. (D) Pathways enriched in the putatively novel pan-cancer core-fitness genes (not belonging to the gene sets listed in B). (E) Gene families enriched in the putatively novel pan-cancer core-fitness genes (not belonging to the gene sets listed in B).

**Table S5:** Pan-cancer and cancer-specific ANOVA results associating gene essentialities to cancer driver events.

**Table S6:** Target priority scores used to compute the significance threshold for identifying high priority targets.

**Table S7:** Final target priority scores across tissues and at the pan-cancer level with detailed partial scores, annotations and formulas to recompute them if changing the score weights (only for tissue specific priority scores).

**Table S8:** Differential dependency markers combined across tissues.

**Table S9:** Genome-wide small-molecule and antibody tractability annotations with information about approved/pre-clinical drugs and their indications.

**Table S10:** PANTHER annotations and protein classes' enrichments of high priority targets, across tractability groups.

**Table S11:** Differential expression and gene set enrichment analysis results comparing the basal expression profiles of cell lines where GPX4 is a significant fitness genes versus those of the other cell lines.

**Table S12 :** sgRNA used for WRN validation experiments.

**Table S13** : Cell lines used for WRN validation experiments.
